# Structure of a Putative Terminal Amidation Domain in Natural Product Biosynthesis

**DOI:** 10.1101/2024.10.28.620694

**Authors:** Michael R. Rankin, Dheeraj Khare, Lena Gerwick, David H. Sherman, William H. Gerwick, Janet L. Smith

## Abstract

Bacteria are rich sources of pharmaceutically valuable natural products, many crafted by modular polyketide synthases (PKS) and non-ribosomal peptide synthetases (NRPS). PKS and NRPS systems typically contain a thioesterase (TE) to offload a linear or cyclized product from a carrier protein, but alternative chemistry is needed for products with a terminal amide. Several pathways with amidated products also possess an uncharacterized 400-amino acid terminal domain. We present the characterization and structure of this putative terminal amidation domain (TAD). TAD binds NAD with the nicotinamide near an invariant cysteine that is also accessible to an intermediate on a carrier protein, indicating a catalytic role. The TAD structure resembles cyanobacterial acyl-ACP reductase (AAR), which binds NADPH near an analogous catalytic cysteine. Bioinformatic analysis reveals that TADs are broadly distributed across bacterial phyla and often occur at the end of terminal NRPS modules, suggesting many amidated products may yet be discovered.

## INTRODUCTION

Bacteria, fungi, and plants produce secondary metabolites of widely varying structure and bioactivity. Many prescribed therapeutics are natural products, including the powerful antibiotic vancomycin^1^, the chemotherapeutic paclitaxel^2^, and the immunosuppressant cyclosporine^3^. An even larger fraction of drugs in clinical use is derived from or inspired by natural products. The enzymes that produce these amazing molecules are potential biocatalysts for chemical modifications that improve or introduce new bioactivities.

Many complex natural products of the polyketide and nonribosomal peptide (NRP) classes are synthesized by enzyme megasynthases. Among these, the modular polyketide synthases (PKS) and non-ribosomal peptide synthetases (NRPS) function as biosynthetic assembly lines of ‘modules’ that act in a distinct sequence. PKS and NRPS modules are multi-enzyme proteins where acyl or peptidyl intermediates are tethered via thioester bonds to the phosphopantetheine (Ppant) cofactor of an acyl carrier protein (ACP) or peptidyl carrier protein (PCP) domain. The carrier protein shuttles the thiotemplated intermediates to each enzyme within the module before transfer to the next module in the pathway. Following the final step of biosynthesis, an enzyme in the terminal module offloads the pathway product.

Typically, a thioesterase (TE) domain releases the pathway product by hydrolysis to a free carboxylic acid or cyclization to a macrolactone or macrolactam. Most NRPS and PKS products are released by a TE located at the C-terminus of the terminal module of the pathway^4–6^. However, for those linear natural products containing a terminal amide and not a carboxylate, an alternative to TE release must exist. A mechanism for amidative release was established for myxothiazol biosynthesis^7^, wherein glycine is condensed with the terminal intermediate through a typical NRPS extension reaction. An embedded monooxygenase then catalyzes the cleavage of the glycine Cα-N bond, yielding a mature amide product and glyoxyl-PCP^7^. Sarpeptins A-B^8^ and zwittermicin A^9^ are formed in a similar manner.

Other NRP natural products contain terminal amides, for example carmabins A-B^10^, vatiamides E-F^11^, and hectoramide B^12^ (**Figure 1**) from marine cyanobacteria, but the corresponding biosynthetic gene clusters (BGCs) do not encode enzymes associated with the glycine-dependent amidation mechanism. Instead, these pathways possess an uncharacterized, 400-amino acid domain at the C-terminus of the terminal NRPS module. The terminal domains of the CarI, VatR and HcaD polypeptides, annotated as proteins with a predicted amino acid dehydrogenase domain, have highly similar sequences (85-95% pairwise sequence identity).

**Figure 1.**
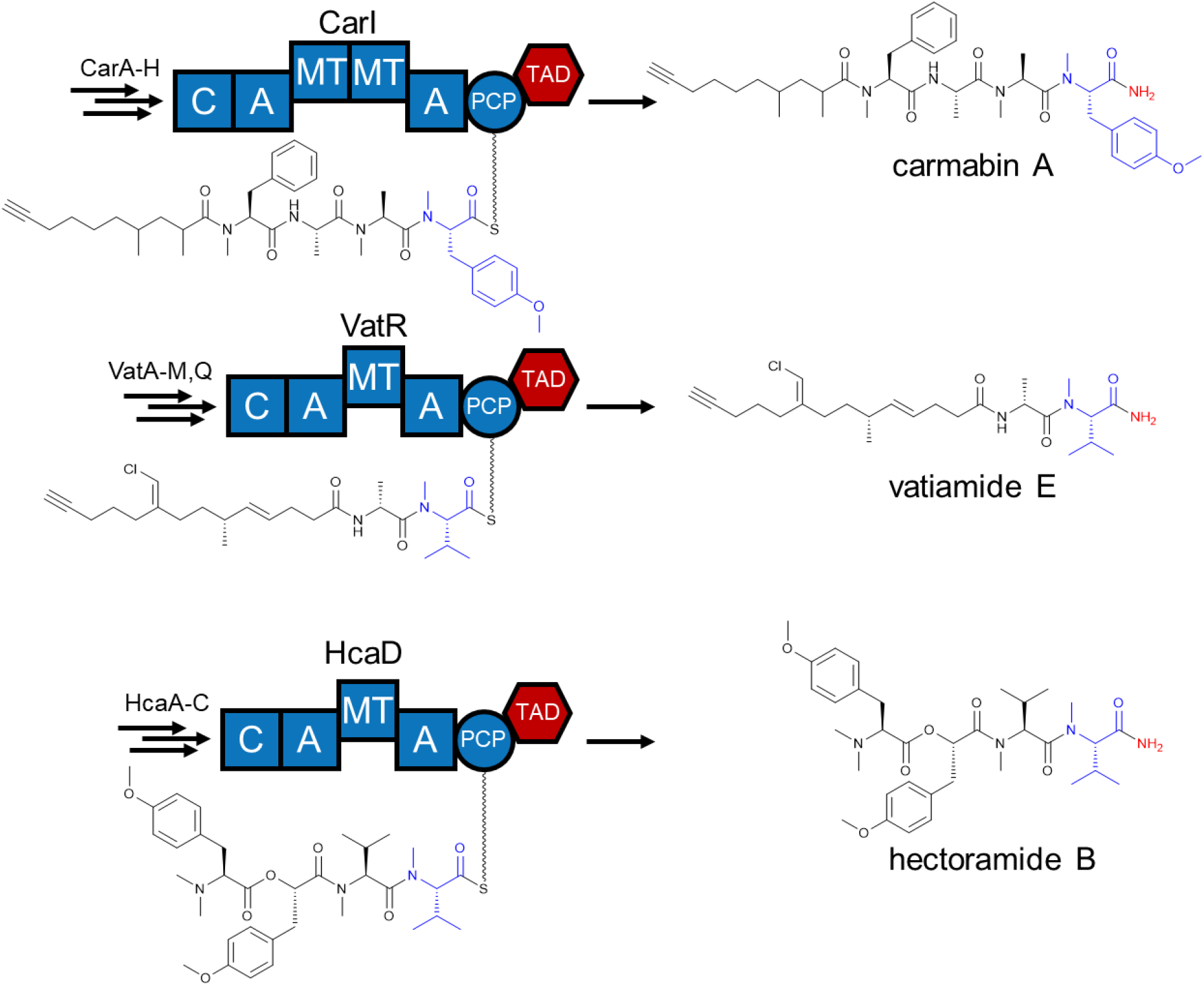
Final modules of carmabin A^17,18^ (genome accession number: GCA_001942475.1), vatiamide E^11^ (GCA_010671925.1), and hectoramide B^12^ (GCA_001854205.2) biosynthesis. Each TAD may release terminally amidated products. Blue atoms and bonds indicate the monomer installed on the current module, while the red atoms may be installed by TAD activity. C: condensation; A: adenylation; PCP: peptidyl carrier protein; MT: methyltransferase.

Hypothesizing that this domain is responsible for both thioester offloading and amide formation, we refer to the domain as a terminal amidation domain (TAD). Other natural products with terminal amides – dragonamides A-E^13^, dragomabin^14^, majusculamides A-B^15^, and almiramides A-C^16^ (**Figure S1**) – are produced by related marine cyanobacteria. Their biosynthetic gene clusters are unidentified, but a TAD is likely. Understanding the mechanism and requirements of the TAD may allow TE-TAD substitution, creating new products with alternative termini and new biological activities.

Here, we report the discovery of broad TAD distribution in the bacterial kingdom. Biochemical characterization and crystal structures for the TAD from the terminal module (CarI) of the carmabin biosynthetic pathway reveal a binding site for NAD(H) with the nicotinamide adjacent to an invariant cysteine residue. Crosslinking studies further show that the cysteine is accessible to substrates tethered to the CarI PCP. The TAD has substantial similarity to a cyanobacterial acyl-ACP reductase. In combination, the results allow the development of possible reaction schemes.

## RESULTS

### Discovery of TAD homologs

We first sought to assess the biological distribution of TAD homologs and to identify any homologs with established function. We designed an approach that was unlimited by prior assumptions, independent of protein context, and initiated with a BLAST^19^ search for CarI TAD homologs in the RefSeq non-redundant protein sequence database. We then used an in-house script to extract the TAD-aligned region of each sequence, excluding duplicates and sequences that aligned with only short regions of TAD. The resulting 757 unique sequences were used to construct a sequence similarity network (SSN)^20,21^ of TAD homologs. Sequence pairs with alignment score > 90 (∼65% sequence identity) were connected, resulting in five major clusters that included 85% of the sequences (**Figure 2A**). We discovered that TAD homologs are broadly distributed across bacterial phyla (**Figures S2-S7**), far beyond the few marine cyanobacterial producers of natural products with established structures and BGCs (**Figure 1**). None of the identified homologs is from a eukaryotic source, and we identified none with an established function. Most of the TAD homologs are encoded in ORFs within genomic (complete or partial) or metagenomic sequences. Based on the flanking sequences, most homologs in four of the five largest SSN clusters are located at the C-terminus of an NRPS protein, strongly suggestive of a natural product termination function. Cluster 4 is an exception, consisting of discrete proteins that do not appear to be BGC products. The consensus sequences for Clusters 1-5 have 24-42% pairwise identity, excepting Clusters 1 and 5 with 59% identical consensus sequences (**Figure S8**). We also used the ESI-GDN (Genome Neighborhood Tool)^20,21^ to search for a potential nitrogen-generating partner for the TAD homologs, but identified no consistently co-occurring candidate ORFs within ten genes of the TAD coding sequence.

**Figure 2.**
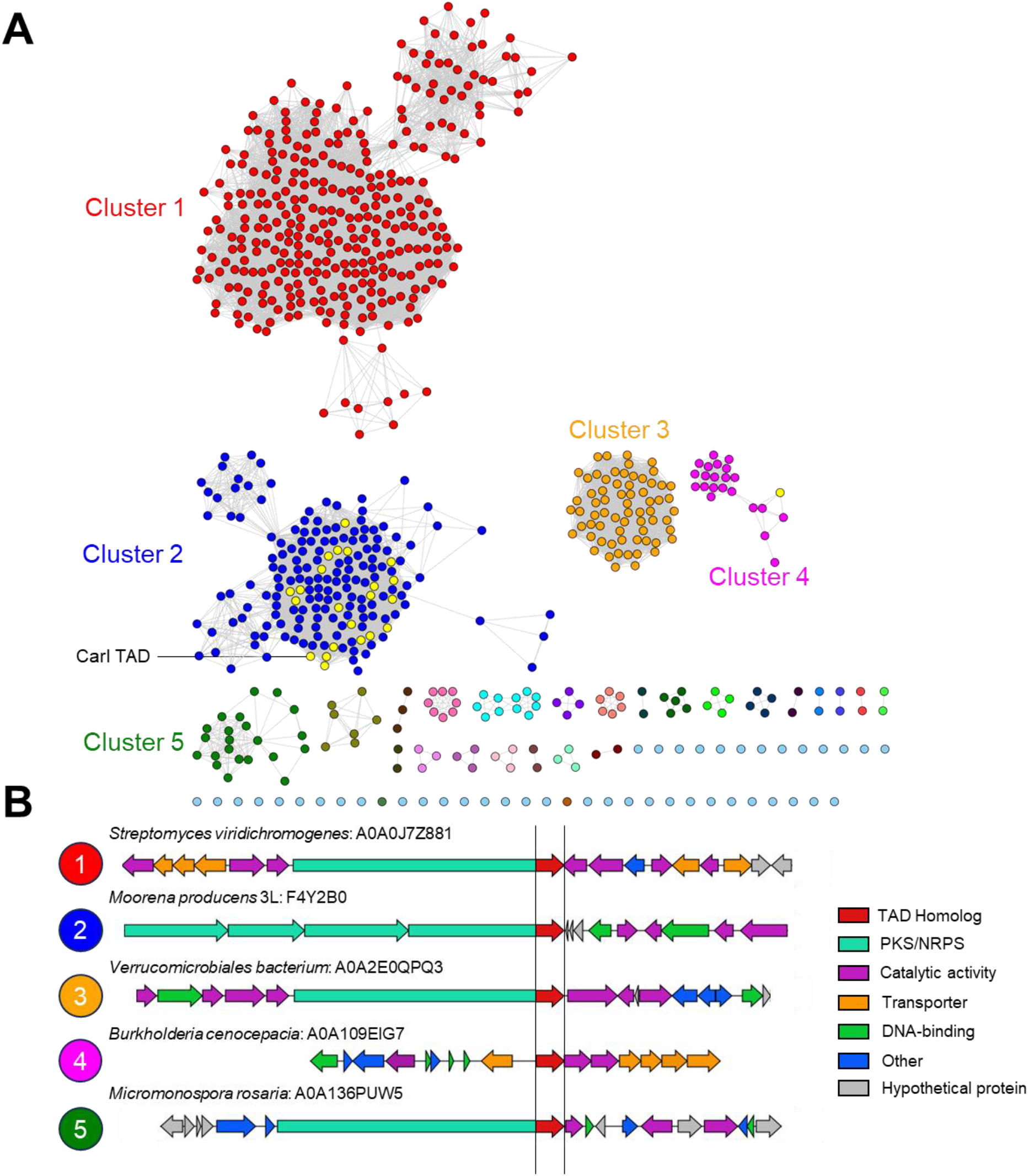
CarI TAD homologs identified in diverse bacteria. **A)** Sequence similarity network (SSN)^20,21^ for proteins related to CarI TAD. Lines connect sequence nodes with high identity (approx. 65%). Clusters defined in this manner are separately colored, with cyanobacterial sequences highlighted in yellow. **B)** Genome neighborhoods for representative examples of the five largest clusters. Many TAD homologs are at the C-terminus of an NRPS module, but those in Cluster 4 are discrete proteins. UniProt IDs are displayed for each TAD homolog; the CarI TAD represents cluster 2.

Our analysis identified several cyanobacterial BGCs containing more than three PKS or NRPS modules that end with a TAD (**Figure S9**). Many of these are hybrid PKS/NRPS pathways that contain aspartate β-hydroxylase domains^22^, suggesting potential siderophore function. Interestingly, a terminal two-module NRPS fragment was identified from *Moorena* sp. SIO3A2. This ORF encodes the machinery to generate the final two amino acids of almiramide A^16^, suggesting a genetic origin for this natural product (**Figure S9A**).

### TAD crystal structure

For a deeper understanding of the TAD, we solved a crystal structure of the TAD excised from CarI using selenomethionyl (SeMet) protein (**Figure 3**). The TAD is a dimer in solution and in crystals **(Figure S10)**. The monomer consists of distinct N- and C-terminal domains (**Figure S11**). The N-terminal domain (NTD, residues 1908-2043, secondary structures β1-β6 and α1-α6) has a topology similar to the macrodomain family^23^, which may bind ADP-ribose and its derivatives^24^. We tested ADP-ribose binding in TAD crystals, but observed no binding to the NTD. The NTD also forms the TAD dimer interface through contacts of helix α1 and the helix α4 C-terminus (**Figure S10C**).

**Figure 3.**
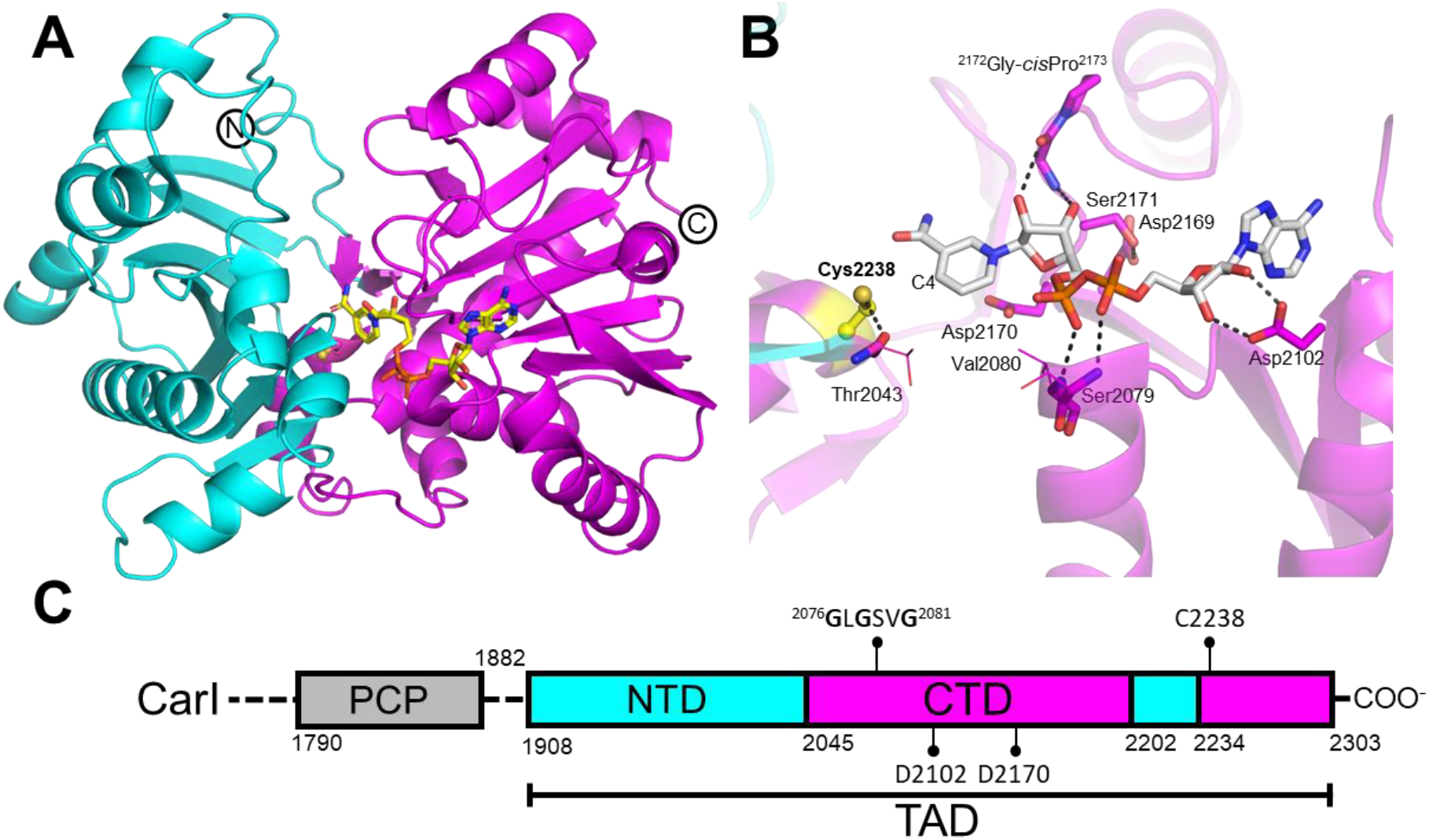
Crystal structure of the CarI TAD. **A)** TAD monomer with cyan NTD and magenta CTD. NAD is bound in a cleft between the two lobes, near putative catalytic Cys2238. **B)** Details NAD binding site. Cys2238 is near the hydride-reactive nicotinamide C4 atom. Black dashed lines indicate hydrogen bonds. **C)** Domain schematic for the TAD region of CarI. The PCP is separated from the TAD by a 26-residue linker. The CTD is interrupted by a 32-residue excursion into the NTD.

The TAD C-terminal domain (CTD, residues 2044-2201 and 2234-2302, secondary structures β7-β13, β15-β18, α7-α10, η1 (3_10_), and α12-α14) has the Rossmann fold common to NAD(P)H-dependent enzymes. Thirty-two amino acids (2202-2233) from the CTD cross over into the NTD where they contribute β14 and α11 (**Figure S11**). The CTD contains a canonical ^2076^GlyxGlyxxGly^2081^ P-loop motif^25^ for binding the NAD(H) diphosphate. Conserved Asp2102 makes the site selective for NAD(H) over NADP(H). Hydrogen bonds of the Asp2102 side chain to both NAD adenosine hydroxyl groups (O2’ and O3’) effectively block binding of both NADP(H) (2’-phosphate) and CoA (3’-phosphate) (**Figure 3B**). We used a thermal shift assay to evaluate potential TAD ligands and detected binding of NAD(H), dephospho-CoA and ADP-ribose, but not NADP(H) or CoA (**Figure S12**).

Among proteins in the structure database, a cyanobacterial acyl-ACP reductase (AAR)^26^ bears similarity to the two-domain TAD structure (**Figure S13**). AAR produces a free aldehyde from an acyl-ACP^26–28^. It is not part of a multi-domain protein, does not catalyze nitrogen transfer, and was not among the hits identified with our initial BLAST search, apparently due to the low (9%) sequence identity of the AAR NTD to the CarI TAD NTD (**Figure S14**). Although the folding topologies are identical, the TAD and AAR NTDs have striking differences. The AAR NTD does not form a dimer, lacks the first β-strand and α-helix of the TAD, has no secondary structures inserted from the CTD, and binds an oxidase that acts on the AAR aldehyde product^26^. The AAR uses a double-displacement “ping-pong” mechanism, first transferring the acyl chain from an acyl-ACP to a catalytic cysteine and releasing holo-ACP (Ppant-ACP), then catalyzing NADPH-dependent reduction of the acyl-enzyme thioester and release of the free aldehyde^27,28^. The CTDs of TAD and AAR have similar structures (**Figure S13**) and sequences (17% sequence identity, **Figure S14**) and bind their respective NAD(P) cofactors similarly (**Figure S13**).

Several conserved amino acids surround the nicotinamide in the TAD NAD complex (**Figures 3B, S15**). Cys2238, located at the beginning of helix α12, is invariant among TAD homologs (**Figure S14**). This amino acid is spatially equivalent to the AAR catalytic cysteine (Cys294) that forms the acyl-enzyme intermediate^26^. Similar to NADP binding to the AAR, NAD binds TAD through a tunnel leading to Cys2238. In the TAD-NAD complex, the Cys2238 thiol forms a hydrogen bond with the Thr2043 carbonyl at the NTD-CTD junction (**Figure 3B**). An invariant ^2042^Thr-Thr-Gly^2044^ motif (**Figure S14**) enables this interaction: the Gly2044 Cα is less than 5 Å from the Cys2238 Cα, and the 2043-2044 peptide is positioned by a hydrogen bond of the Thr2042 hydroxyl to the Gly2044 NH. The nicotinamide C4 (position of hydride transfer) is 4 Å from the Cys2238 thiol, and N1 is near invariant Asp2170 (**Figure 3B**). The interaction of the negatively charged Asp2170 carboxylate may favor the binding of the positive NAD^+^ nitrogen over the neutral NADH. Interestingly, members of cluster 3, which are mostly comprised of Verrucomicrobiota, contain an invariant tyrosine rather than an aspartate (**Figure S15**). In AAR, a glycine occupies the analogous position. Asp2170 is part of a conserved ^2169^Asp-Asp-Ser-Gly-*cis*Pro^2173^ loop. The loop structure is stabilized by the *cis* configuration of Pro2173 and by a number of hydrogen bonds, including the NH and C=O of Gly2172 with the nicotinamide ribose hydroxy groups (**Figure 3B**). In contrast, the well-ordered Asp2170 side chain forms no hydrogen bonds, but is nearly stacked with the nicotinamide ring (<3.5 Å to C6). Like Asp2170 and despite the conserved surroundings, the well-ordered nicotinamide also forms no hydrogen bonds with the protein, precluding assignment of the amide flip. The amide group is not co-planar with the aromatic ring, tilted by ∼30° (**Figure S16B**).

### NAD and phosphopantetheine binding in a conserved cleft

The conserved NAD site in TAD together with published assays of the AAR highlighting the role of a catalytic cysteine^26–28^ strongly suggest a mechanism in which the first step is substrate transfer from peptidyl-PCP to Cys2238. Thus, we initially loaded the PCPs of CarI and VatR with a substrate mimic (CarI: hexanoyl-dimethyltyrosyl; VatR: succinyl-methylvalyl). In independent reactions, the CarI and VatR TADs were incubated with or without combinations of NAD(H) and various metals, under a range of pH conditions. Ammonia and all 20 amino acids were tested as potential nitrogen donors. Release of substrate from the PCP was monitored by coupled liquid chromatography/mass spectrometry (LC/MS), but substrate release was not detected under any of the tested conditions, indicating an inadequate substrate or a missing component. Without access to authentic substrates (**Figure 1**), we undertook alternative characterization approaches. Assuming a substrate is delivered by the PCP of the module containing the TAD, we tested whether the Ppant-PCP can reach Cys2238.

We used the thiol-reactive crosslinker bismaleimidoethane (BMOE)^29^ to probe a PCP-TAD interaction. When the CarI holo-PCP and TAD were incubated *in trans* in the presence of BMOE, a link of one PCP and one TAD was detected by denaturing polyacrylamide gel electrophoresis (SDS-PAGE) as a dominant band with migration matching the untreated, fused PCP-TAD didomain. Intact protein MS data confirmed this crosslink (**Figure S17-S18**). Peptide mapping tandem mass spectrometry (MS/MS) identified Cys2238 as a crosslinking partner (**Figure 4** and **S19**). An additional crosslink was detected between the PCP Ppant thiol and TAD Cys2258, a non-conserved amino acid located on the TAD surface. No reproducible crosslinks to the other six cysteine residues in the TAD were detected. We compared the relative reactivity of Cys2238 and Cys2258 with the PCP Ppant using C2238A, C2258S and C2238A/C2258S substitutions in CarI TAD. The double substitution prevented nearly all crosslinking, the C2238A substitution led to a major decrease (25% crosslinked) and the C2258S substitution to a moderate decrease (70% crosslinked) (**Figure S20**). This supports the idea that Cys2238 is the preferred target of the PCP Ppant, even though it is less accessible than Cys2258 (12.5 Å^2^ vs 44.2 Å^2^ exposed surface area).

**Figure 4.**
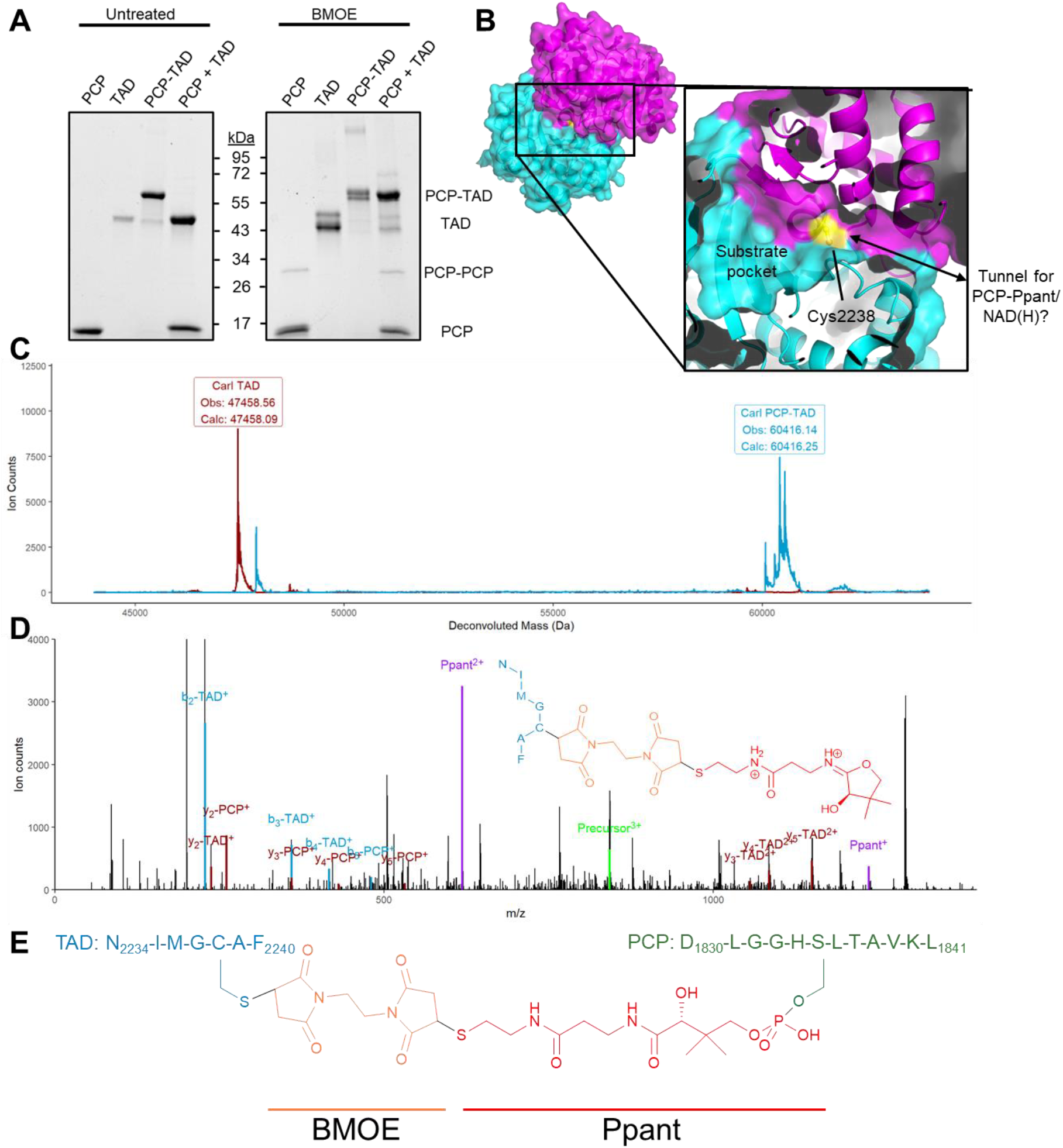
Crosslinking of the PCP Ppant and TAD Cys2238. **A)** SDS-PAGE gel of CarI PCP, TAD, PCP-TAD didomain, or PCP and TAD proteins incubated with and without BMOE. **B)** Cys2238 central location in the cleft between TAD NTD (cyan) and CTD (magenta). This residue is adjacent to a pocket that may accommodate the waiting substrate and the tunnel (arrow) where NAD(H) binds. **C)** Deconvoluted mass spectrum of intact CarI PCP and TAD proteins following incubation with (blue) and without (red) BMOE. Cysteine, added to quench the crosslinking reaction, was detected as a cap to several BMOE modification sites. See **Figure S18** for details. **D)** Detection of a crosslinked peptide consisting of TAD residues 2234-2240 and Ppant-PCP 1830-1841 linked by BMOE. The MS/MS spectrum includes a Ppant ejection fragment^30^ linked to the TAD peptide containing Cys2238. The first isotopic peaks for important ions are highlighted. See **Figure S20** for details. **E)** Chemical structure of the crosslink between chymotryptic fragments of TAD Cys2238 and the Ppant-PCP thiol.

Cys2238 is located in a wide cleft between the TAD NTD and CTD, and is also at the innermost end of the tunnel where NAD binds. To test whether the tunnel is a common access route for the PCP Ppant and NAD(H), crosslinking experiments between the PCP and TAD C2258S were performed in the presence and absence of NADH (**Figure 5A**). The addition of NADH decreased the extent of PCP-TAD crosslinking, suggesting that NAD binding blocks Ppant access, *i*.*e*. Ppant enters through the NAD-binding tunnel. The TAD had moderate affinity for both NAD^+^ (K_d_ = 11.6 µM) and NADH (K_d_ = 3.5 µM), with slight preference for NADH (**Figure S12B**). The binding is sufficiently weak to allow cofactor diffusion out of the shared binding site, creating space for an incoming Ppant-substrate. To visualize Ppant in this site, we cocrystallized CarI TAD with 3’-dephosphocoenzyme A (dPCoA), a Ppant analog. The dPCoA is bound in the tunnel similarly to NAD. The Ppant reached Cys2238 at the end of the NADH-binding tunnel and formed a disulfide bond during crystallization (**Figure 5B**). This result demonstrates that Ppant can access and react with the Cys2238 thiol. In the disulfide form, the Cys2238 side chain was rotated towards the site occupied by the nicotinamide C4 and amide in the NAD complex, a ∼90° rotation relative to the Cys2238 position when hydrogen bonded to the Thr2043 carbonyl.

**Figure 5.**
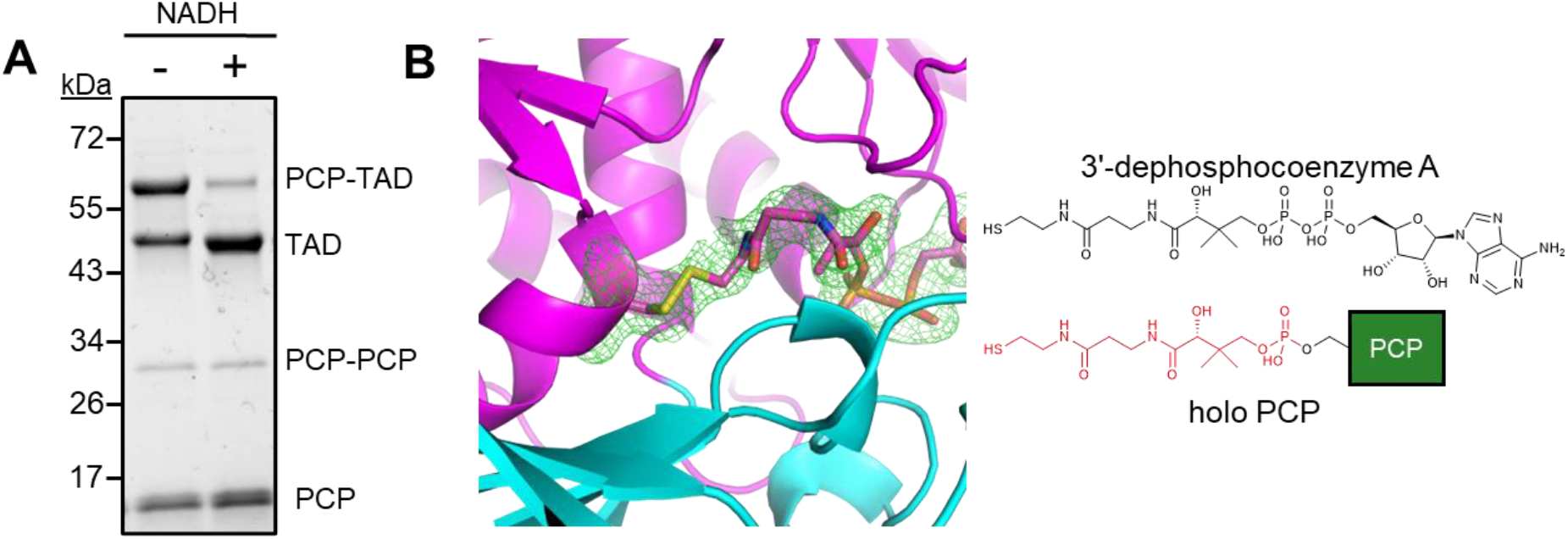
Competing Ppant and NAD interaction with TAD. **A)** BMOE crosslinking of CarI PCP and CarI TAD C2258S. NADH prevented PCP-Ppant access to Cys2238. **B)** Co-crystal structure of Ppant mimic dPCoA with the CarI TAD. The Ppant formed a disulfide with Cys2238 (Polder density^31^ contoured at 3σ).

## DISCUSSION

Through sequence similarity analysis, we found evidence of broad TAD distribution throughout bacterial phyla, well beyond the three marine cyanobacterial strains identified prior to this study^11,12,17^. The TAD consists of two independently evolved subdomains: an NTD with a macrodomain fold and a dehydrogenase-like, nucleotide-binding CTD that is specific for NAD(H) over NADP(H) or CoA. The conservation of the fused didomain suggests a common overall function for these gene products. This aspect of our study highlights the importance of unbiased searches into the vast encoded proteome to identify uncharacterized protein families and unearth clues about function. Our approach differs from previous studies in constructing an SSN from the query sequence (here TAD) using only the region within each “hit” that aligns to the query. Given the propensity for DNA sharing among bacteria and the existence of large, multi-functional proteins in natural product biosynthesis, this approach is critical for discovery of family relationships among domains.

The TAD family clearly shares a common ancestor with AAR, although the TAD and AAR NTDs have diverged substantially in sequence (9% identity) and function. Their more similar CTDs (17% identity) suggest a partially shared mechanism, including a critical role for an invariant cysteine (CarI Cys2238) in the cleft between the NTD and CTD. Our crosslinking results and the sequence conservation suggest a substrate access route to Cys2238 through the NAD(H) tunnel. These findings led us to consider potential reaction schemes, beginning with transfer of the substrate from the PCP to TAD Cys2238 and release of holo-PCP in a step identical to AAR^26^ (**Figure 6**).

**Figure 6.**
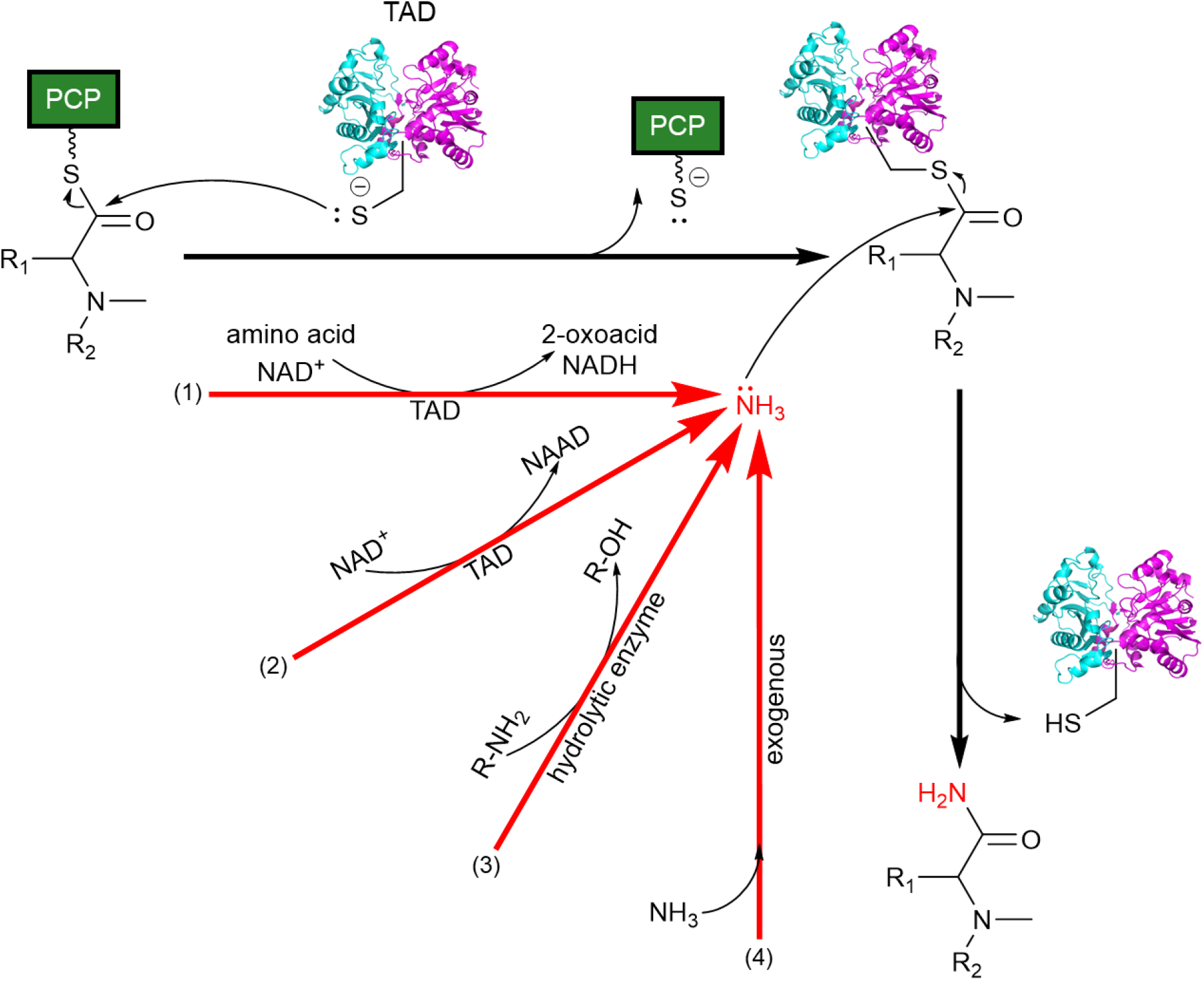
Proposed reaction schemes for TAD activity. After transthioesterification of the substrate to the TAD Cys2238, a terminal amide may be formed by ammonia attack on the peptidyl-*S*-enzyme thioester. Ammonia may be generated by (1) NAD^+^-dependent oxidation of an amino acid, (2) salvaged from the NAD amide, (3) produced by an unknown hydrolytic enzyme, or (4) supplied exogenously. (R_1_: side chain of the C-terminal amino acid of the natural product intermediate, R_2_: remainder of the intermediate, wavy line: Ppant).

Although the source of the amide nitrogen remains unknown, we reason that nucleophilic attack by ammonia on a TAD thioester intermediate is most plausible. We envision four sources of ammonia. First, the TAD may oxidize the α-carbon of an amino acid, forming ammonia and the corresponding 2-oxoacid. This scheme requires NAD and an appropriate amino acid. Many amino acid oxidizing enzymes are known, although none is a TAD homolog. Second, the TAD may hydrolyze the NAD amide, yielding ammonia and nicotinic acid dinucleotide (NAAD). Amidohydrolases that convert nicotinamide to nicotinic acid or nicotinamide mononucleotide (NMN) to nicotinic acid mononucleotide (NaMN)^32^ have been characterized, although these are not TAD homologs. Interestingly, a few TAD-encoding BGCs had an NAD synthetase gene immediately downstream of the cluster (**Figure S9**). NAD synthetase converts NAAD to NAD. Third, TAD could partner with an external enzyme that releases ammonia directly to the thioester intermediate. We searched the carmabin, vatiamide, and hectoramide biosynthetic gene clusters and proximal genomic regions for genes whose products may produce ammonia and partner with the TAD. Due to ammonia lability, such partner enzymes would need tight coupling to the TAD thioester intermediate. One such example is the glutaminase subunit/domain of a glutamine amidotransferase^33^. However, no candidates were identified that were shared by a number of BGCs. Finally, nucleophilic attack by exogenous ammonia is possible, as in NH_3_-dependent NAD synthetase^34^, but the nearby conserved cofactor binding site and lack of reaction with ammonia suggests alternative chemistry.

The above hypotheses deviate from the AAR mechanism, as the NAD cofactor is not used for thioester reduction. However, another route to amide formation may involve the reduction of the thioester to an aldehyde, followed by nitrogen incorporation and oxidation back to an amide. This NADH-dependent, yet redox-neutral mechanism, designated a complex NAD-dependent transformation^35^, is used by 1-L-*myo*-inositol-1-phosphate synthase^36^, uridine diphosphate (UDP)-galactose-4-epimerase^37^, and ornithine cyclodeaminase^38^.

As more biosynthetic pathways and associated products are discovered, new chemoenzymatic strategies are identified. The TAD reported here is one such example, but there is still much to learn about how cells terminate PKS/NRPS pathways by amidation. For example, the NRPS module that produces bovienimides A and B lacks an obvious termination mechanism, but yields a linear amide (bovienimide A) or acid (bovienimide B)^39^. Termination may be accomplished by a yet undiscovered enzyme, or initiated by the nucleophilic attack of environmental ammonia or water on the PCP thioester to furnish the final product^39^.

Decades of BGC sequencing have revealed considerable rearrangement of coding sequences within and between organisms and BGCs – representing Nature’s experiments in expanding biosynthetic capacity. When analyzing and comparing functional regions, they must be considered not only in the context of full-length multi-functional enzymes where they exist, but also individually together with homologs in different protein contexts.

## CONCLUDING REMARKS

With each newly characterized PKS/NRPS domain, our biocatalytic toolbox grows. Here, we present an enigmatic domain that appears to terminate and amidate the product of a hybrid PKS/NRPS cluster. Sequence analysis reveals that these domains are widespread in bacteria, with 757 unique homologs identified. The structure of the domain is highly reminiscent of cyanobacterial acyl-ACP reductase, which provided great insight into a potential mechanism.

With this knowledge, a series of mass spectrometric experiments were performed to identify a central cysteine residue that is likely critical for TAD activity, as well as the tunnel through which the substrate-carrying Ppant enters. This study lays a foundation for further exploration of TAD enzyme activity. Once the minimal requirements for TAD function are determined, thioesterase-TAD substitutions could be made to alter the final product of a PKS/NRPS cluster. In the case of a thioesterase domain with cyclization activity, a TAD would terminate the pathway with a linear amide instead of a macrocycle, yielding a product with the potentially vastly different bioactivity.

## QUANTIFICATION AND STATISTICAL ANALYSIS

### Crystallographic data collection and processing

Statistics for crystallographic data collection and processing, structure determination, and refinement were calculated with the phenix.table_one tool from the Phenix package v1.20.1-4487.66. These statistics are summarized in **Table S1**.

## ACKNOWLEDGMENTS

This work was supported by NIH grant R01 DK042303 and the Rita Willis Professorship to JLS, by NIH grant R35 GM118101 and the Hans W. Vahlteich Professorship to DHS, by R01 GM107550 to LG and WHG, and by NIH grants T32 GM145304 and F31 CA265082 and a Life Sciences Institute Cubed grant to MRR. We thank Lydia Freddolino (University of Michigan) for the script to isolate the TAD homology regions for SSN generation, and Wendy Feng (University of Michigan Life Sciences Institute Mass Spectrometry Core) for assistance with MS analysis.

Eli Eisman cloned the CarI TAD enzyme. GM/CA@APS has been funded by the National Cancer Institute (ACB-12002) and the National Institute of General Medical Sciences (AGM-12006, P30GM138396). This research used resources of the Advanced Photon Source, a U.S. Department of Energy (DOE) Office of Science User Facility operated for the DOE Office of Science by Argonne National Laboratory under Contract No. DE-AC02-06CH11357.

## AUTHOR CONTRIBUTIONS

M.R.R. and J.L.S. designed the research. M.R.R. and D.K. performed the research. M.R.R., D.K., and J.L.S. analyzed the data. M.R.R. and J.L.S. wrote the paper with input from all authors.

## METHOD DETAILS

### SSN generation

The EFI-EST^1,2^ server generates SSNs from the full protein sequences returned in a BLAST search or from domains defined by a specified Pfam or InterPro family. The TAD is not a member of any defined family, and many hits from the TAD query were large, multidomain proteins. In order to create an SSN from only the TAD region, a BLAST^3^ search of the RefSeq non-redundant (nr) protein sequence database was performed using CarI residues 1908-2303 as the query, yielding 1844 hits. A script (Lydia Freddolino, U Michigan) was used to truncate hit sequences to include only those regions that aligned to the query (**Figure S21**). This script also pruned sequences that did not align with at least 300 residues of CarI TAD. 950 sequences remained after this pruning step. For clustering with the EFI-EST server, an alignment score of 90 was chosen to connect nodes, corresponding to approximately 65% sequence identity.

Finally, redundant sequences were consolidated into a single, representative node, resulting in a total of 757 unique sequences of TAD homologs. BGCs were rendered using the CAGECAT^4^ server clinker tool^5^.

### Molecular cloning

All polymerase chain reaction (PCR) amplifications were performed using the KOD Hot Start DNA Polymerase (MilliporeSigma) and the primers listed in **Table S2**. The *carI* gene (UniProt ID: F4Y2B0) was subcloned into the pET29b(+) vector (Novagen) by restriction digestion using NdeI and XhoI enzymes (New England Biolabs). Cosmid DNA containing a fragment of the carmabin biosynthetic cluster (accession number: HQ696503.1) was used as a template. The resulting plasmid and same cloning strategy were used to create CarI TAD and CarI PCP expression constructs.

Constructs encoding CarI TAD Δhelix and CarI PCP-TAD were generated by creating ligation-independent cloning (LIC) inserts compatible with pMCSG7^6^ and pMocr^7^, respectively (**Table S3**). These inserts were processed with T4 DNA polymerase (MilliporeSigma) and assembled into their destination vector. Site-directed mutagenesis was performed using the QuikChange II XL Site-Directed Mutagenesis Kit (Agilent). All constructs were confirmed by sequencing.

### Protein production and purification

*Gene Expression*. Expression plasmids were transformed into *E. coli* BL21 (DE3) pRare2-CDF^*8*^ cells that were made competent by the Mix & Go! *E. coli* Transformation Kit (Zymo Research). Terrific broth (TB) cultures containing 50 µg mL^-1^ kanamycin (or 100 µg mL^-1^ ampicillin for pMCSG7 or pMocr vectors) and 50 µg mL^-1^ spectinomycin were grown at 37 °C with shaking at 225 rpm until an OD_600_ of 1.0 was reached. Cultures were cooled to 20 °C for 1 hr, then induced with 200 µM IPTG (isopropyl β-D-1-thiogalactopyranoside) and 2 g L^-1^ of L-arabinose, grown 18 hr, and harvested by centrifugation at 12,000 x g.

To produce PCP-containing proteins in the holo state, plasmids were transformed into the *E. coli* BL21 (DE3) BAP1 cell line^9^, which constitutively expresses *sfp*, encoding a nonspecific phosphopantetheinyl transferase^10^. Conversely, to produce apo-form carrier proteins, expression of *E. coli entD* was repressed by addition of a trace metal cocktail to the culture media^11^.

*Protein Purification*. Cell pellets were resuspended in 70 mL lysis buffer per liter of culture (50 mM HEPES pH 7.8, 300 mM NaCl, 10% (v/v) glycerol, 20 mM imidazole pH 7.8), augmented with 1 mg mL^-1^ chicken lysozyme (Sigma), 50 µg mL^-1^ bovine DNase I (Sigma), and 2 mM MgCl_2_, then incubated 30 min at room temperature with agitation. Complete lysis was achieved via sonication (Branson Sonifier 450). Following centrifugation at 30,000 x g, the soluble fraction was collected, filtered, incubated 2 hr with 5 mL of packed Ni-NTA agarose beads (Qiagen), filtered (0.45 µm Millex-HP PES membrane filter unit, Millipore) and loaded onto a glass chromatography column (Bio-Rad). The beads were washed with 100 mL of lysis buffer before the protein was eluted in 40 mL elution buffer (50 mM HEPES pH 7.8, 300 mM NaCl, 10% (v/v) glycerol, 400 mM imidazole pH 7.8).

When purifying the CarI TAD Δhelix or PCP-TAD proteins, the His_6_ or His_6_-Mocr^7^ tag was cleaved with tobacco etch virus (TEV) protease by incubation with 1 mg TEV protease/ 40 mg protein, supplemented with 5 mM DTT. The mixture was dialyzed (Spectra/Por dialysis membrane, 3.5 kDa molecular weight cutoff (MWCO), Spectrum) overnight against 2 L gel-filtration buffer (50 mM HEPES pH 7.8, 150 mM NaCl, 10% (v/v) glycerol) with 2 mM DTT. To separate the target protein from uncleaved protein and TEV protease, the mixture was applied to a 5 mL HisTrap HP column (GE Healthcare) at a flow rate of 2.5 mL min^-1^, followed by 5 column volumes of lysis buffer. The target protein was collected in the flowthrough.

For a final purification step by gel filtration, proteins were concentrated to 5 mL using a centrifugal filter unit (Amicon) with an appropriate MWCO, and injected onto either a Superdex 75 or Superdex 200 Hiload 16/60 prep grade gel-filtration column (GE Healthcare), pre-equilibrated with gel filtration buffer. Eluates were assessed by SDS-PAGE. Target fractions were pooled, concentrated, flash-frozen in liquid nitrogen, and stored at -80 °C.

CarI hexanoyl-*N,O*-dimethyltyrosyl-PCP and VatR succinyl-*N*-methylvalyl-PCP substrates for TAD activity assays were prepared by purifying the regions of CarI and VatR that contain condensation, adenylation, and methyltransferase domains. These purified proteins were then incubated with ATP, SAM, and the respective amino acid (CarI: Tyr, VatR: Val). After PCP loading was complete, the PCPs from the preceding module, which had been acylated by Sfp as above (CarH: hexanoyl, VatQ: succinyl), were added to the reaction. PCP modification was assessed using intact protein or peptide mapping MS.

*Selenomethionine Protein Production*. SeMet CarI TAD protein was produced using the protocol above but with the following alterations. The CarI TAD plasmid was transformed into E. coli BL21 (DE3) cells, and 60 mL starter cultures were grown overnight in Luria Broth (LB) media containing 50 µg mL^-1^ kanamycin overnight at 37 °C with agitation at 225 rpm. The culture was harvested at 3,000 x g, then washed once with SeMet Medium (Molecular Dimensions) to remove residual methionine. After resuspension with 60 mL SeMet Medium, 5 L cultures of SeMet Medium supplemented with 50 µg mL^-1^ kanamycin and 5 mL L^-1^ SeMet Solution (Molecular Dimensions) were inoculated and grown to an OD_600_ of 1.4 before being induced with 200 µM IPTG. The purification continues as above.

### Protein crystallization and structure determination

Initial CarI TAD crystals (residues 1888-2303) were grown in 24-well sitting-drop vapor diffusion trays with 250 µL of reservoir solution. Crystals of the free enzyme were obtained when 1.1 µL 10.9 mg mL^-1^ protein was mixed with 0.9 µL mother liquor consisting of 100 mM Tris-HCl pH 8.5, 200 mM MgCl_2_, 28.4% PEG 3350, and 5.33% 1,1,1,3,3,3-hexafluoro-2-propanol. To obtain SeMet derivative crystals, 1 µL CarI TAD (SeMet) at 10 mg mL^-1^ was mixed with 1 µL mother liquor (100 mM Tris-HCl pH 8.5, 150 mM MgCl_2_, 26% PEG 3350, 5% 1,1,1,3,3,3-hexafluoro-2-propanol) and grown as the wild type. For NADH-bound crystals, 3 mM NADH was added to a 10 mg mL^-1^ solution of CarI TAD protein before mixing 1 µL protein with 1 µL reservoir solution (100 mM Tris-HCl pH 8.5, 200 mM MgCl_2_, 28.4% PEG 3350, 3% 1,1,1,3,3,3-hexafluoro-2-propanol, and 3% 1,6-hexanediol). Each condition yielded hexagonal plates overnight at 20 °C.

The CarI TAD (1888-2303) crystal structure revealed an N-terminal helix that was poorly ordered in the crystal lattice, presumably part of the linker between the PCP and TAD (**Figure S22**). Given that residues 1888-1907 seemed detrimental to crystal packing, we made a shortened CarI TAD Δhelix (residues 1908-2303), which crystallized in a new space group with greatly improved data quality.

CarI TAD Δhelix (residues 1908-2303) crystals were grown in 24-well sitting-drop vapor diffusion trays at 4 °C with a reservoir volume of 250 µL. 0.9 µL protein at 6.6 mg mL^-1^ was added to 0.7 µL reservoir solution (100 mM bis-tris propane pH 8.5, 134 mM sodium iodide, 21.6% PEG 3350). Clusters of hexagonal plates appeared after a week. Crystals of the NAD^+^ complex were grown by adding 0.8 µL 6.6 mg mL^-1^ protein supplemented with 2 mM NAD^+^ to 0.8 µL reservoir solution (100 mM bis-tris propane pH 8.5, 100 mM sodium iodide, 21% PEG 3350, 3% 2-methyl-2,4-pentanediol (MPD)) in a 96-well sitting-drop vapor diffusion tray with a reservoir volume of 50 µL. Butterfly wing-like crystals grew after a week at 4 °C. Crystals of the dPCoA complex were obtained by adding 0.8 µL 6.7 mg mL^-1^ CarI TAD Δhelix protein containing 2 mM dPCoA (Cayman Chemical) to 0.4 µL 100 mM bis-tris propane pH 8.5, 75 mM sodium iodide, and 22% PEG 3350 in a 96-well 3-drop sitting drop vapor diffusion tray held at 4 °C. Butterfly wing-like crystals grew after a week.

All crystals were harvested and flash-frozen in liquid nitrogen without further cryoprotection. X-ray diffraction data were collected as described in **Table S1** and was processed and scaled using XDS^12^. SAD phasing of the long (residues 1908-2303) CarI TAD data was carried out via AutoSol^13^ in the PHENIX package^14^. An initial polyalanine trace was built using Buccanneer^15^, followed by manual model building in *Coot*^16^. Models were refined by phenix.refine^17^. Subsequent CarI TAD datasets were solved by molecular replacement using Phaser^18^ with the current highest-resolution structure as a search model. All structure figures were created using PyMOL^19^. Structure validation was done with MolProbity^20^. Multiple sequence alignments were created using the Clustal Omega^21^ web service in Jalview^22^. The SeAAR alignment to TAD homologs was manually adjusted after comparing the structures of CarI TAD and SeAAR. Exposed surface area calculations were performed with the GETAREA program^23^.

### Thermal shift assay

10 µL samples containing 0.4 mg mL^-1^ CarI TAD, 10 mM cofactor, and 10X SYPRO Orange dye (Invitrogen) in gel-filtration buffer were arranged in a 384-well PCR plate. A QuantStudio 7 Pro Real-time PCR system (Applied Biosystems) was used to monitor fluorescence at an excitation wavelength of 470 nm and emission wavelength of 570 nm. Samples were heated from 25 to 95 °C at a rate of 0.03 °C/s. The melting temperature (T_m_) was determined by taking the maximum of the first derivative of the fluorescence data with respect to temperature using the Protein Thermal Shift software (Applied Biosystems). For the determination of binding affinities, this experiment was repeated with a serial dilution of cofactor. K_d_s were calculated with the Thermott server^24^.

### Chemical crosslinking

Crosslinking assays between the CarI PCP and TAD were performed by incubating 100 µM holo PCP and 50 µM TAD (or PCP-TAD didomain) in 40 mM phosphate buffer pH 7.0, 5% glycerol. The reaction was initiated by adding 200 µM BMOE (Thermo Scientific) in dimethylsulfoxide (DMSO). The reaction proceeded for 90 min at 25 °C. Unreacted maleimide groups of BMOE were quenched with 20 mM L-cysteine.

### Mass spectrometry

*Intact Protein*. Mass spectrometry was conducted using an Agilent 6545 Q-TOF equipped with an Agilent 1290 HPLC. Protein was diluted to 4 µM with 1% formic acid. 1 µL was injected onto a PLRP-S 300 Å, 2.1 × 50 mm, 3 µm column with a flow rate of 0.2 mL/min. A gradient from 4.75% to 95% acetonitrile in 0.1% formic acid was run over 8 min. Positive-mode mass spectra were collected using a fragmentor energy of 225 V and skimmer voltage of 25 V. Spectrum deconvolution was performed using BioConfirm (Agilent).

*Peptide Mapping Tandem Mass Spectrometry*. 3 µL of each crosslinking reaction was diluted to 40 µL in 100 mM Tris pH 8.0, 10 mM CaCl_2._ This mixture was reduced by the addition of 5 mM TCEP (tris(2-carboxyethyl)phosphine) and incubated at 50 °C for 20 min. After cooling to 25 °C, 15 mM iodoacetamide was added and alkylation proceeded for 15 min in the dark.

Chymotrypsin (Sequencing grade, Roche) was then added in a ratio of 1 µg protease: 20 µg sample protein. Digestion continued for 18 hours at 37 °C and quenched with 1.25 µL formic acid. 10 µL peptide mix was injected onto a 2.1 × 150 mm, 2.7 µm AdvanceBio Peptide Map column (Agilent) with flow rate 0.4 mL/min, column temperature 40 °C using the same HPLC/MS instruments as above with a two-step gradient: 4.75% to 40% over 30 min, then 40% to 100% over 20 min. Three spectra/sec were collected in positive-ion mode with 175 V fragmentor energy and 65 V skimmer voltage. For untargeted MS/MS fragmentation, dominant species were isolated with a width of 4 *m/z*. Collision energy (CE) in volts was set by the equation *CE* = 3.2 ∗ (*m*/*z*)/100 + 3. MS/MS fragmentation data was used to discover and confirm the identity of peptides of interest, and Bioconfirm (Agilent) was used to assign peptide sequences to the resulting spectra.

