## Supplemental Information for "Structure of a Putative Terminal Amidation Domain in Natural Product Biosynthesis"

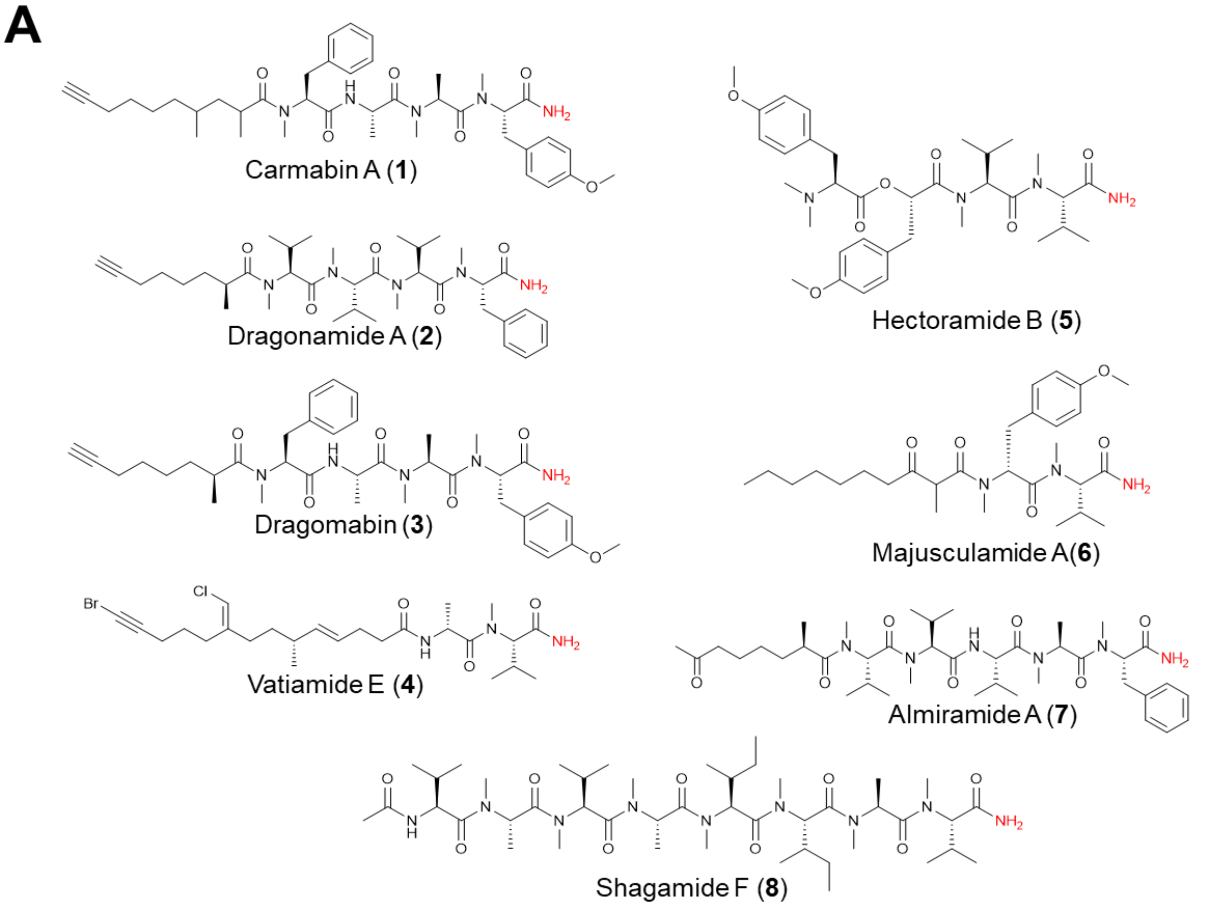


**B**

| Compound(s) | Representative Structure | Producer | TAD? | Pathway accession number |
|---|---|---|---|---|
| Carmabin A[40–42] | 1 | *Moorena producens* 3L | Yes | 5' end: HQ696502.1<br>3' end: HQ696503.1 |
| Dragonamides A-E[43] | 2 | *Moorena producens*[a] | Likely | unknown |
| Dragomabin[44] | 3 | *Moorena producens*[a] | Likely | unknown |
| Vatiamides E-F[45] | 4 | *Moorena producens* ASI | Yes | MK618714.1 |
| Hectoramide B[46] | 5 | *Moorena producens* JHB | Yes | OQ821997.1 |
| Majusculamides A-B[47] | 6 | *Moorena producens* Gomont[a] | Likely | unknown |
| Almiramides A-C[28] | 7 | *Moorena producens* PAB[a] | Likely | unknown |
| Shagamides A-F[48] | 8 | *Inflatella coelosphaeroides* (symbiont?) | Unknown | unknown |

**Figure S1. A)** Chemical structures of select C-terminally amidated molecules of interest. **B)** Information on C-terminally amidated natural products that are or may be produced by TADs. Pathway accession numbers, when available, are indicated. [a]While these original compound isolation papers reported *Lyngbya majuscula* as the producing organism, this genus and species was subsequently revised in the taxonomic literature to *Moorena producens*[25].

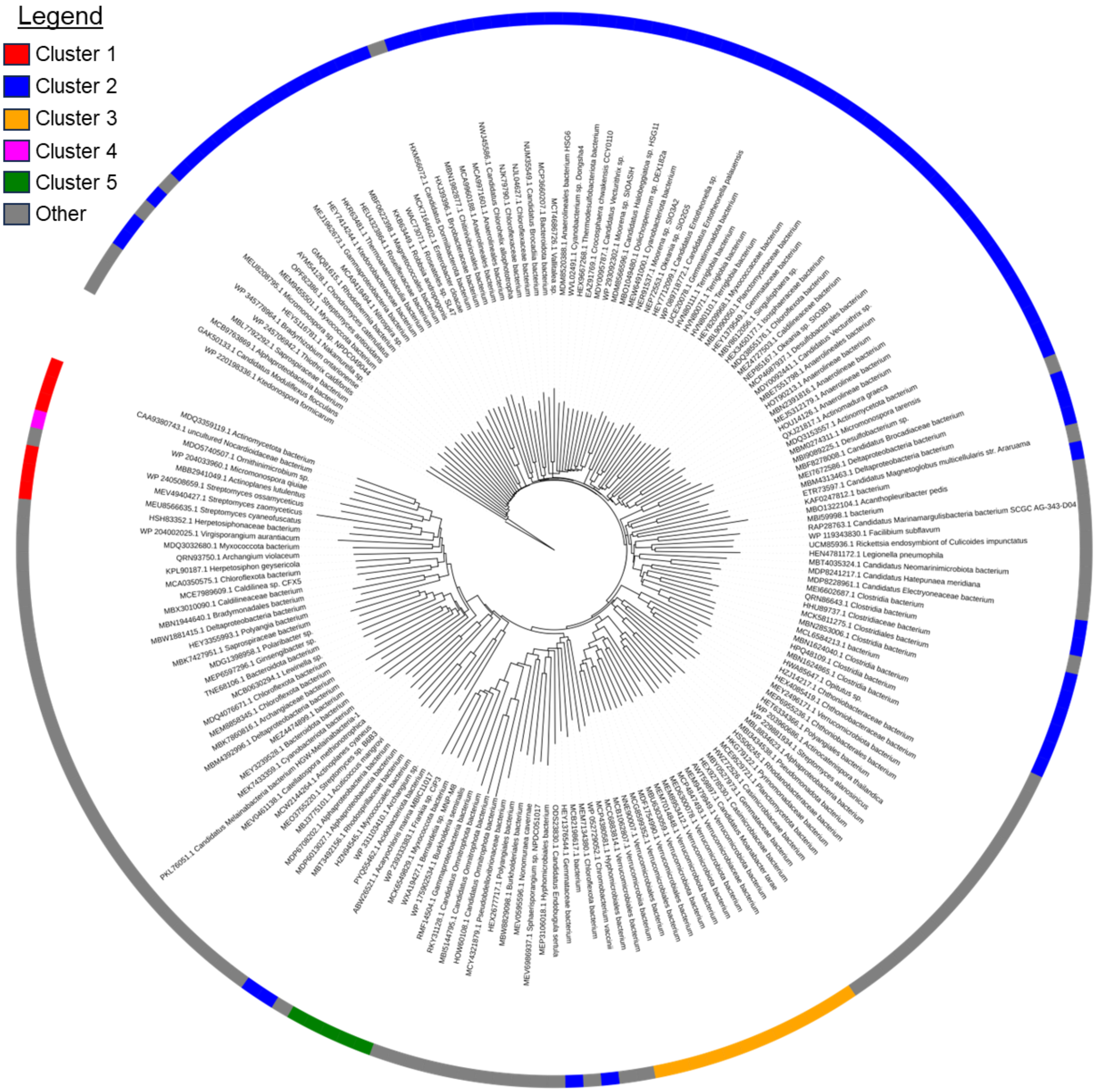


**Figure S2.** Phylogenetic tree illustrating the spread of TAD homologs across bacteria. Protein accession numbers and biological sources are shown on the label of each sequence. A redundancy threshold of 85% was set in Jalview[22], leaving 182 representative sequences. The first five clusters of the SSN are colored (**Figure 2**), with minor clusters and singletons in gray. Trees for each of Clusters 1-5 are in **Figures S3-S7**. This tree was made using iTOL[26].

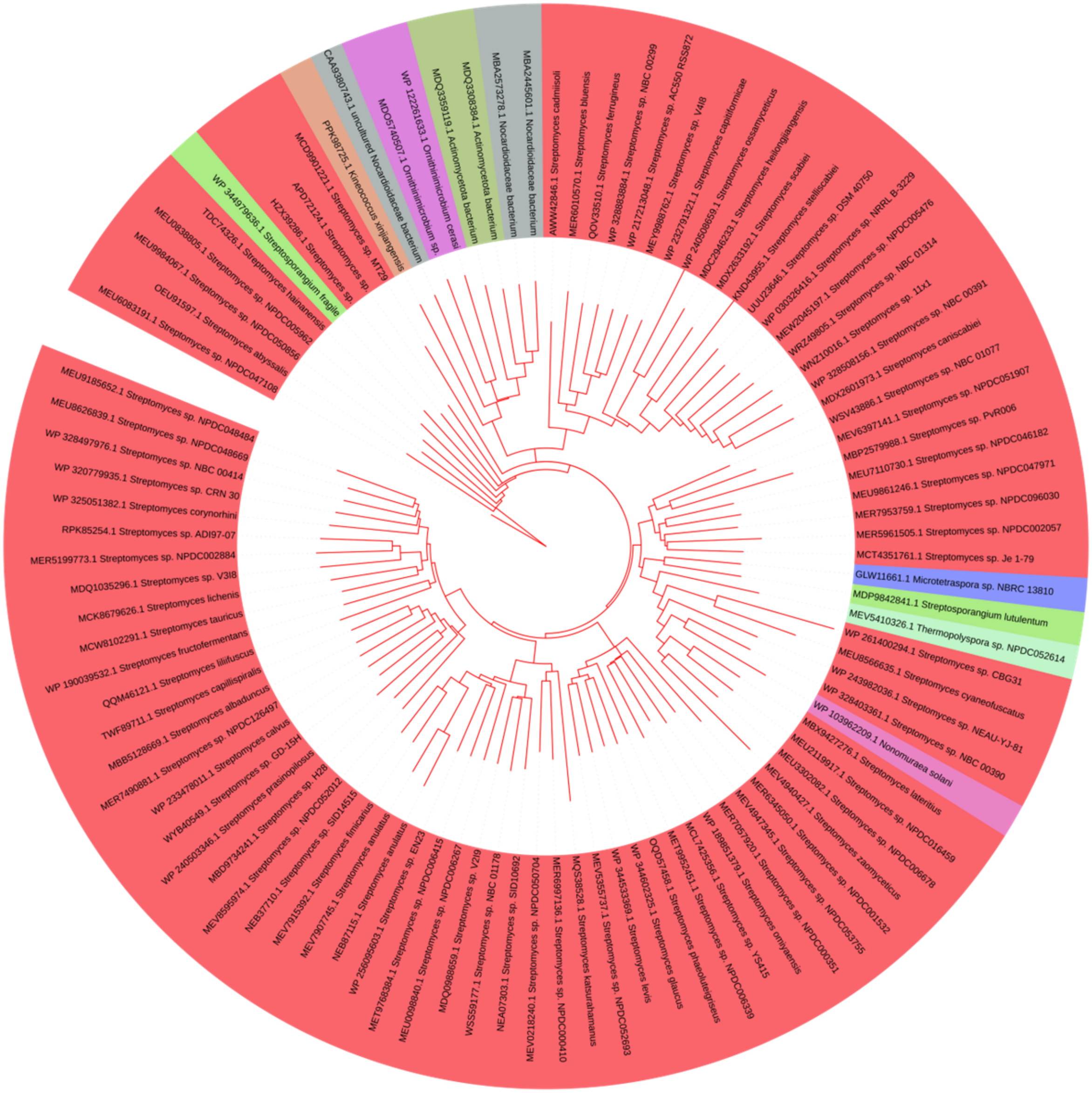


**Figure S3.** Phylogenetic tree of representative sequences from cluster 1 of the CarI TAD SSN (**Figures 2A, S2**). Cluster 1 includes 460 total sequences, 336 unique. Protein accession numbers and biological sources are shown on the label of each sequence. A redundancy threshold of 95% was set in Jalview[22], leaving 98 representative sequences (unique). Sequences are colored by genus. Cluster 1 is dominated by Streptomyces (red). This tree was made using iTOL[26].

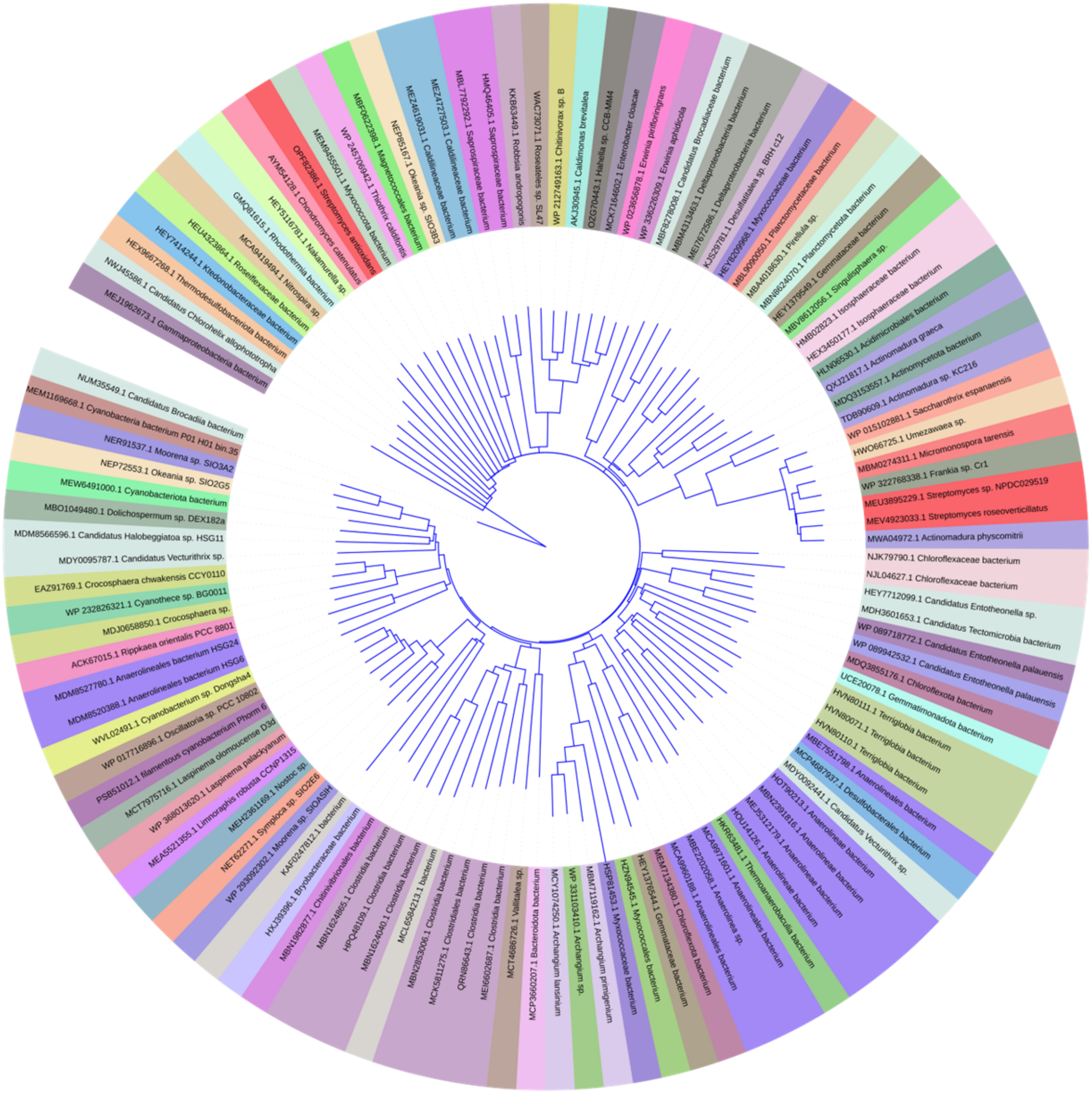


**Figure S4.** Phylogenetic tree of representative sequences from cluster 2 of the CarI TAD SSN (**Figures 2A, S2**). Protein accession numbers and biological sources are shown on the label of each sequence. Cluster 2 includes 207 total sequences, 182 unique. A redundancy threshold of 90% was set in Jalview[22], leaving 114 representative sequences. Sequences are colored by genus. Cluster 2 is the most broadly distributed among bacteria. The CarI TAD is in the group represented by *Moorena sp.* SIO3A2 (blue, just below the open area). This tree was made using iTOL[26].

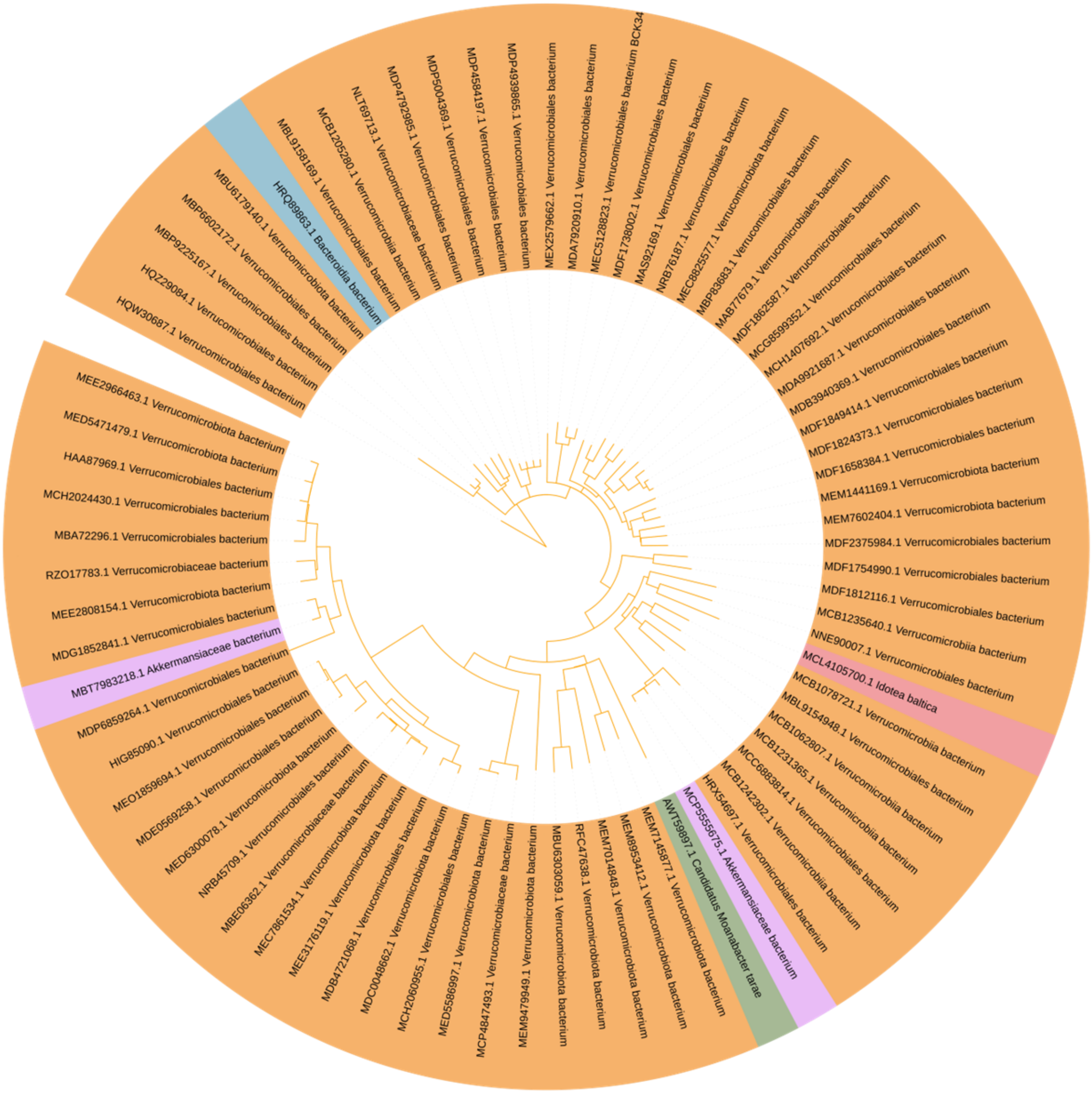


**Figure S5.** Phylogenetic tree of sequences from cluster 3 of the CarI TAD SSN (**Figures 2A, S2**). Protein accession numbers and biological sources are shown on the label of each sequence. Cluster 3 contains 76 sequences (67 unique) and is dominated by Verrucomicrobiota. Sequences are colored by genus. This tree was made using iTOL[26].

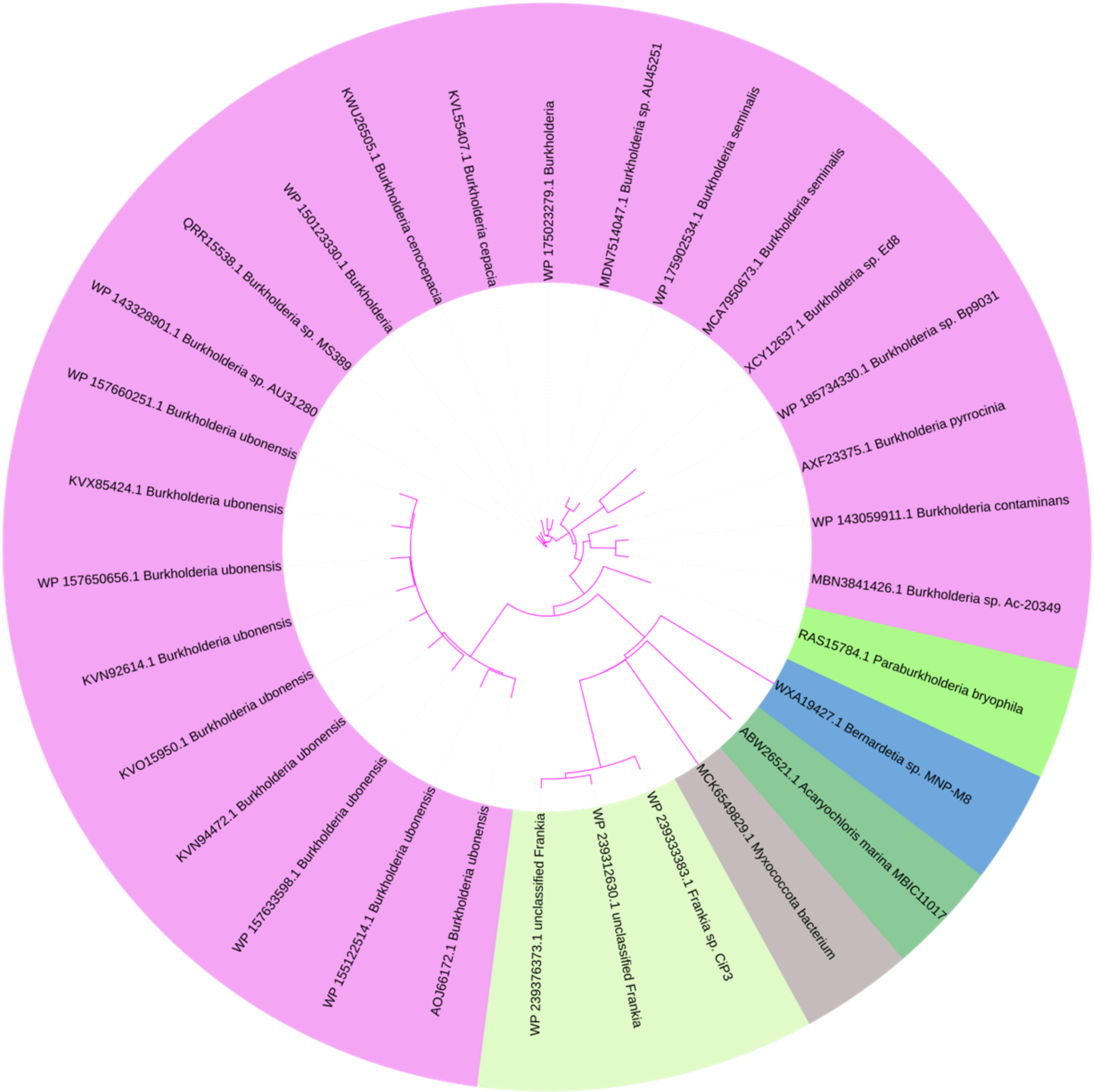


**Figure S6.** Phylogenetic tree of sequences from cluster 4 of the CarI TAD SSN (**Figures 2A, S2**). Protein accession numbers and biological sources are shown on the label of each sequence. Cluster 4 contains 30 sequences (23 unique) and is dominated by Burkholderia. Sequences are colored by genus. This tree was made using iTOL[26].

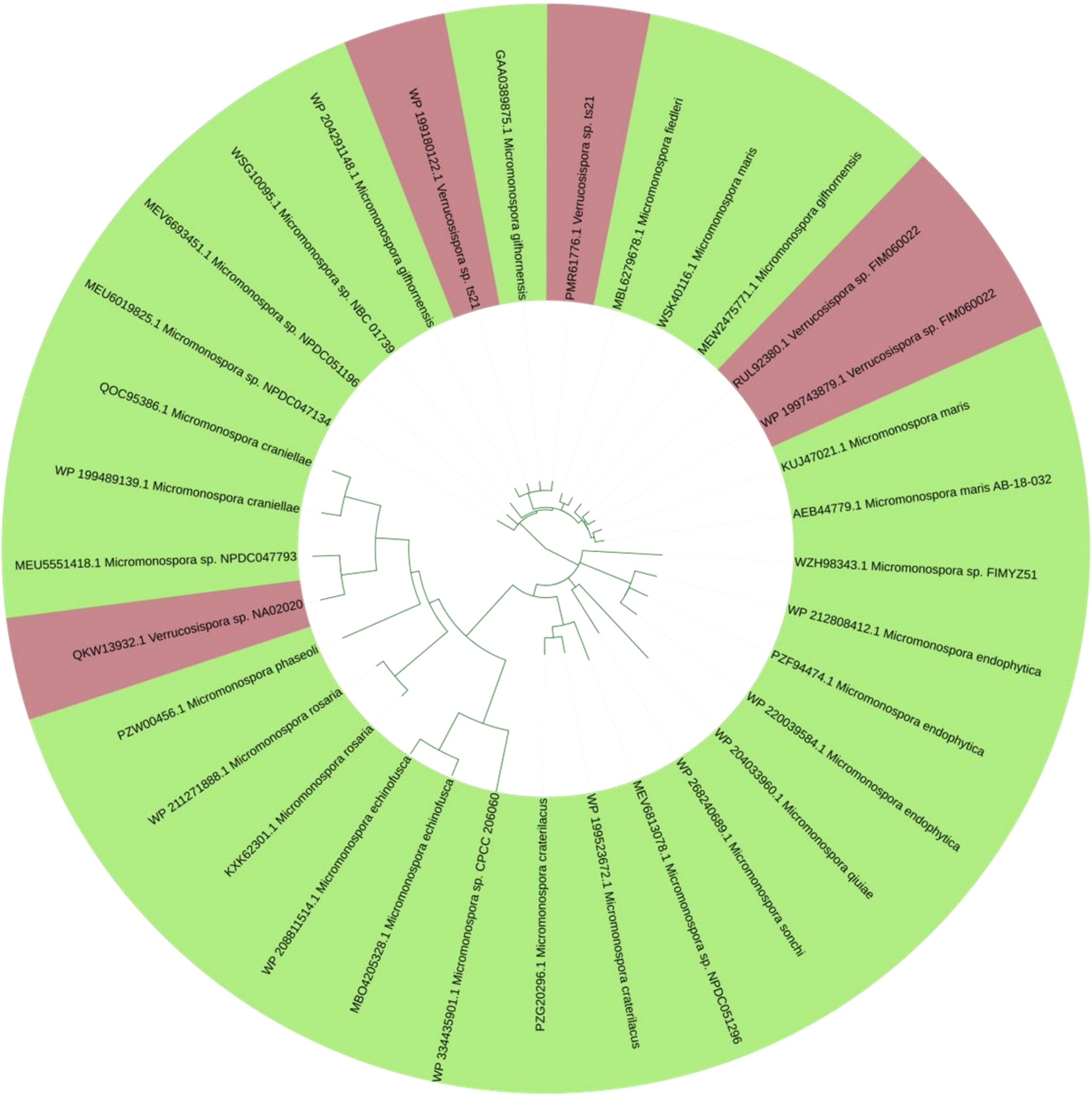


**Figure S7.** Phylogenetic tree of sequences from cluster 5 of the CarI TAD SSN (**Figures 2A, S2**). Protein accession numbers and biological sources are shown on the label of each sequence. Cluster 5 contains 33 sequences (22 unique) and is dominated by Micromonospora. Sequences are colored by genus. This tree was made using iTOL[26].

A

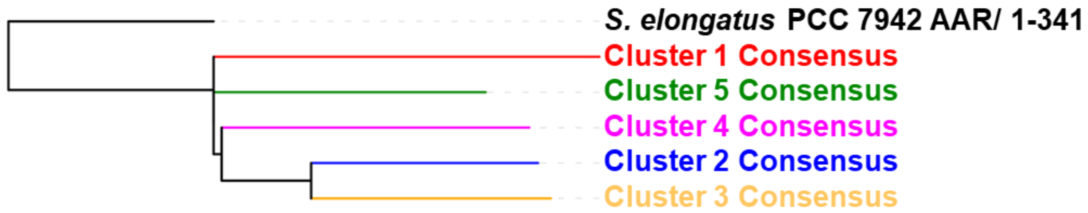


B

| | Cluster 1 | Cluster 2 | Cluster 3 | Cluster 4 | Cluster 5 | AAR |
|---|---|---|---|---|---|---|
| **Cluster 1** | 100 | 34.2 | 30.9 | 30.4 | 58.7 | 25.5 |
| **Cluster 2** | 34.2 | 100 | 42.1 | 26.5 | 33.8 | 20.5 |
| **Cluster 3** | 30.9 | 42.1 | 100 | 24.1 | 32.7 | 17.9 |
| **Cluster 4** | 30.4 | 26.5 | 24.1 | 100 | 31.5 | 21.4 |
| **Cluster 5** | 58.7 | 33.8 | 32.7 | 31.5 | 100 | 26.1 |
| **AAR** | 25.5 | 20.5 | 17.9 | 21.4 | 26.1 | 100 |

**Figure S8.** Relationship between AAR and TAD homologs in five major SSN clusters. **A)** Phylogenetic tree generated with Jalview[22] and visualized in iTOL[26]. **B)** Percent identity matrix for alignment among the five cluster consensus sequences and AAR.

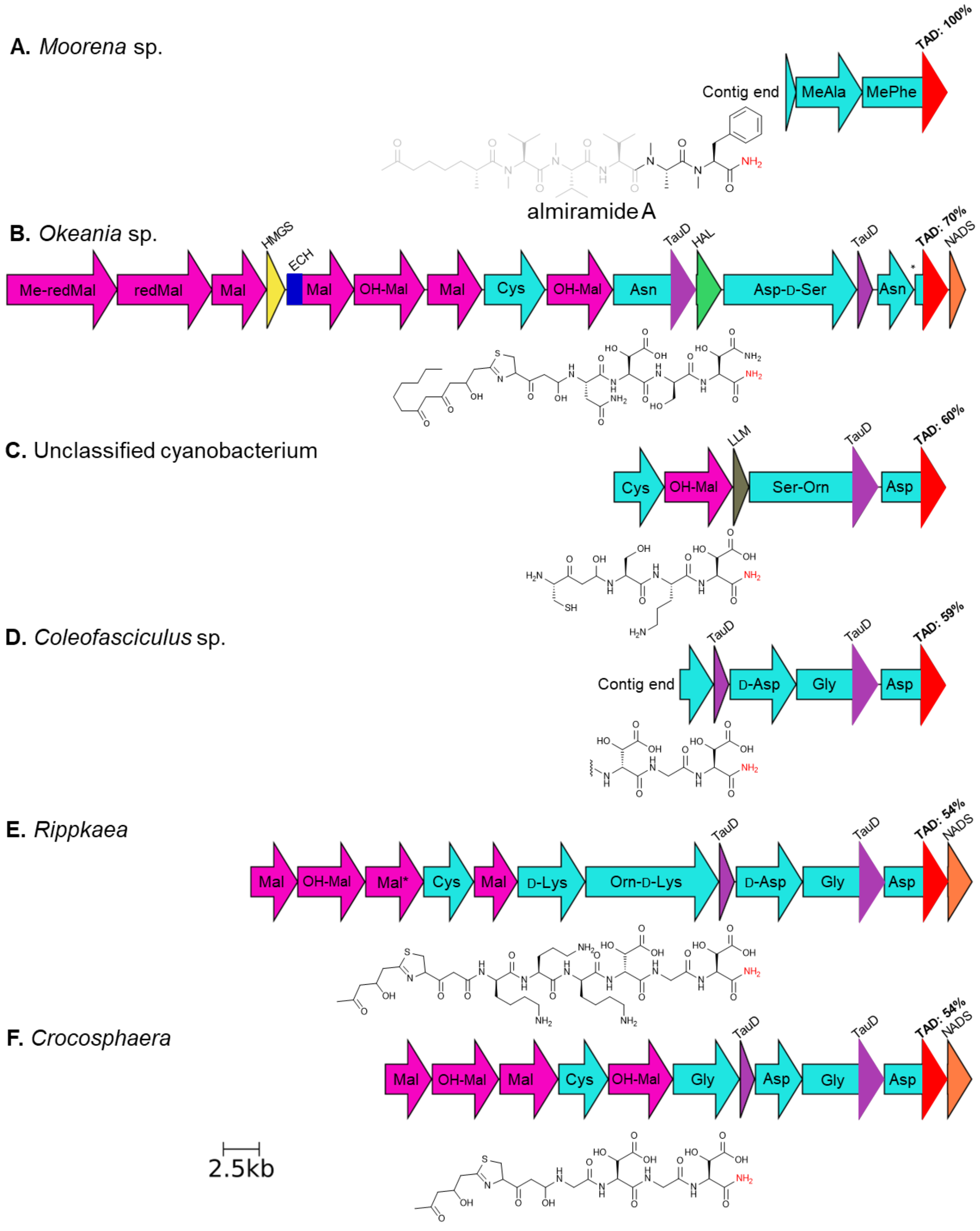


**Figure S9.** Hypothetical natural products produced by cyanobacterial TAD-containing BGCs. The amino acid sequence identity of each TAD (red arrow) relative to CarI TAD is indicated. Each PKS (pink) or NRPS (cyan) polypeptide is labeled with the product(s) predicted by AntiSMASH[27], or the published structure of almiramide[28] with the region not predicted from the identified contig in gray. Mal: malonyl, OH-Mal: hydroxymalonyl, redMal: fully-reduced malonyl, Me-redMal, fully-reduced methylmalonyl, Mal*: PKS with an inactive ketoreductase. Other ORFs are HMGS: 3-hydroxy-3-methylglutaryl synthase[29], ECH: enoyl-CoA hydratase, TauD: aspartate β-hydroxylase[30],

HAL: halogenase, NADS: NAD synthetase, LLM: luciferase-like monooxygenase. An asterisk signifies an inter-module break or sequencing error. Clusters were annotated with AntiSMASH[27] and visualized by the CAGECAT[4] server Clinker tool[5]. Biological sources and accession numbers: **A)** *Moorena* sp. SIO3A2, JAAHHC010000062; **B)** *Okeania* sp. SIO2G4, JAAHGW010000025; **C)** Cyanobacteriota bacterium isolate MAG099, JBFLZA010000003; **D)** *Coleofasciculus* sp. G1-WW12-02, JAVJRX010000111; **E)** *Rippkaea orientalis* PCC 8801, CP001287; **F)** *Crocosphaera chwakensis* CCY0110, AAXW01000011.

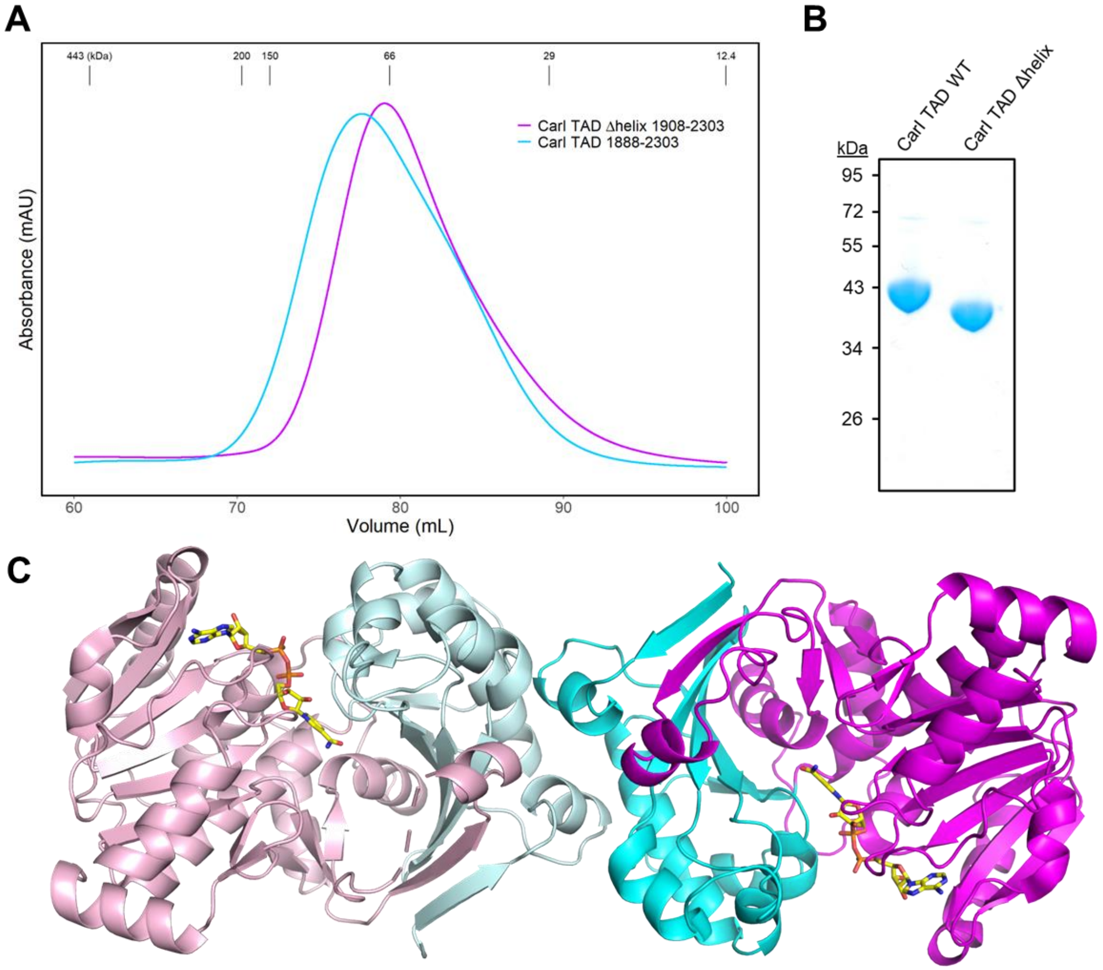


**Figure S10**. Oligomeric state and purity of CarI TAD proteins. **A)** Size-exclusion chromatography indicates that both forms of CarI TAD used in this study are dimeric. CarI TAD 1888-2303 has a mass of 47.5 kDa ,while the TEV protease-cleaved CarI TAD Δhelix 1908-2303 protein is 44.4 kDa. Masses of molecular weight standards are indicated at the top. **B)** SDS-PAGE gel of the same purified proteins. **C)** CarI TAD dimer. The dimer interface is mediated by the NTD (cyans), with the NAD-binding CTD at the dimer periphery (magentas).

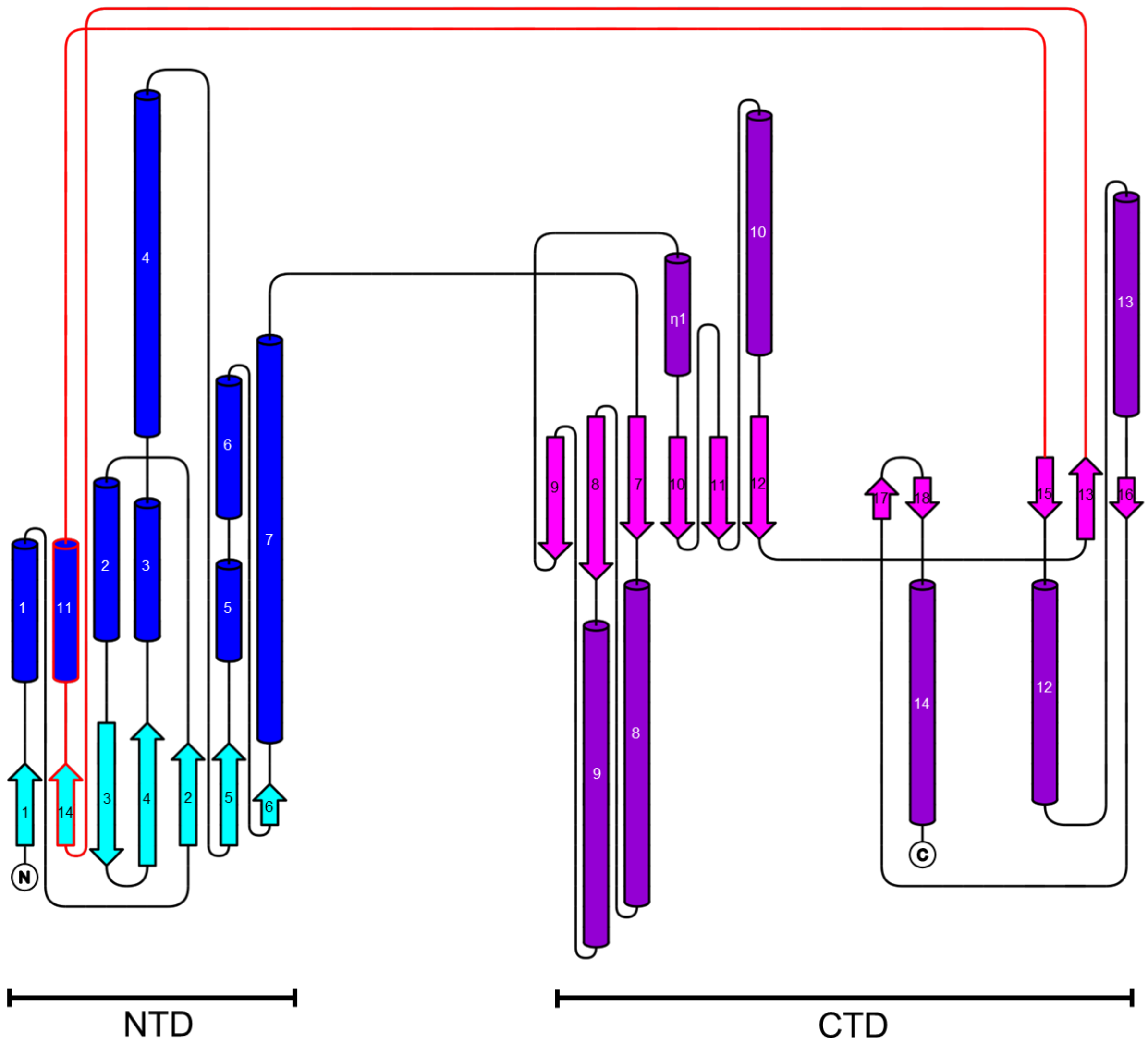


**Figure S11**. Topology diagram of the CarI TAD. Structure topology was determined using PROMOTIF[31] via PDBsum[32] and modified using TopDraw[33]. Secondary structures of the NTD region are in blues, and in magenta/pink for the CTD. The subdomain crossover region is represented by red lines. α-helices and β-strands are numbered individually. One $3_{10}$-helix is marked by an η symbol.

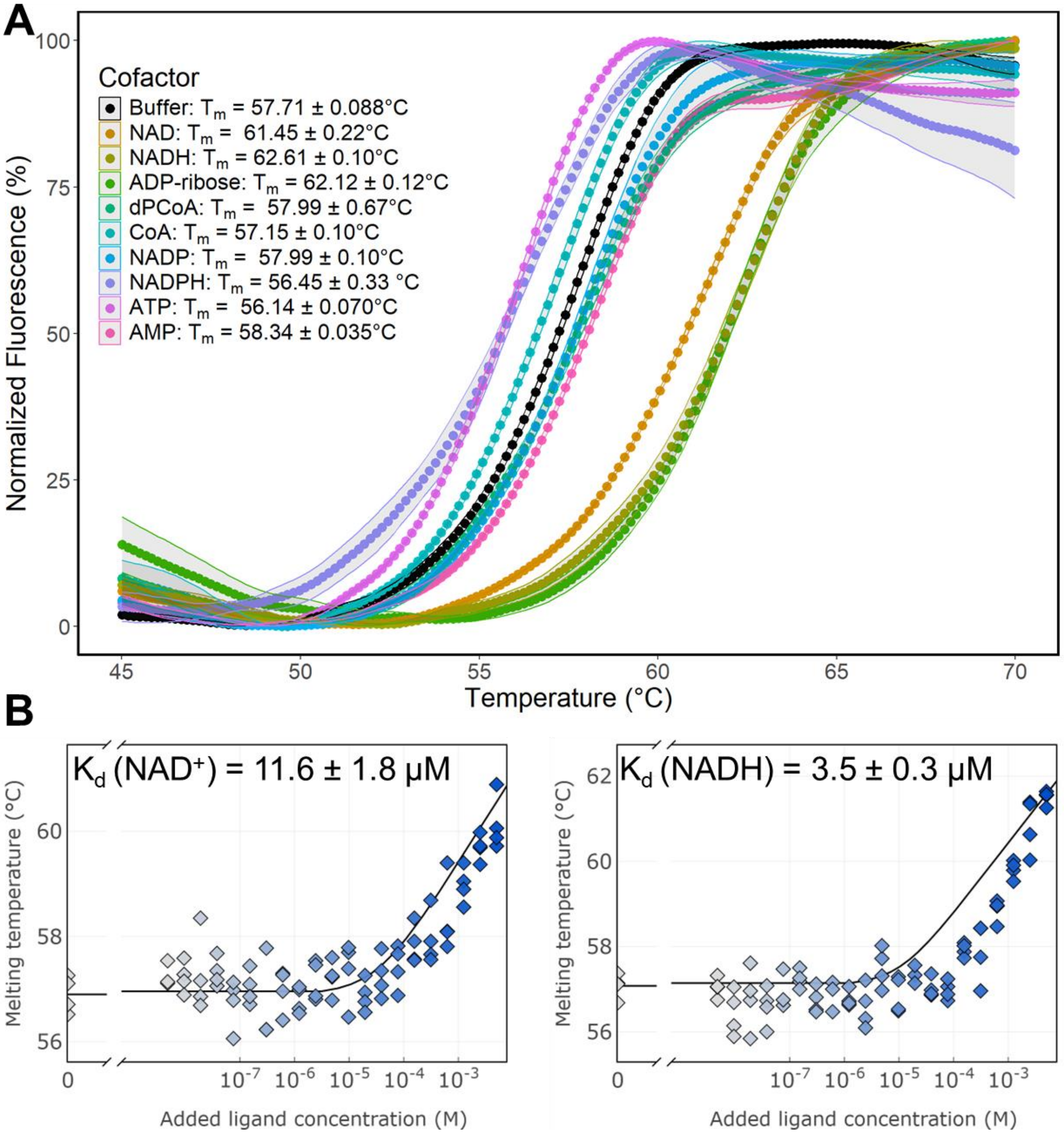


**Figure S12**. Interaction of ligands with the CarI TAD. **A)** Thermal shift assay (TSA) showing the impact of potential ligands on CarI TAD stability. Notably, NAD(H), but not NADP(H), stabilizes the TAD and increases its melting temperature ($T_m$) substantially. Ribbons indicate the standard error of four measurements. **B)** $K_d$ was determined for $NAD^+$ and NADH by TSA. The TAD is selective for NADH.

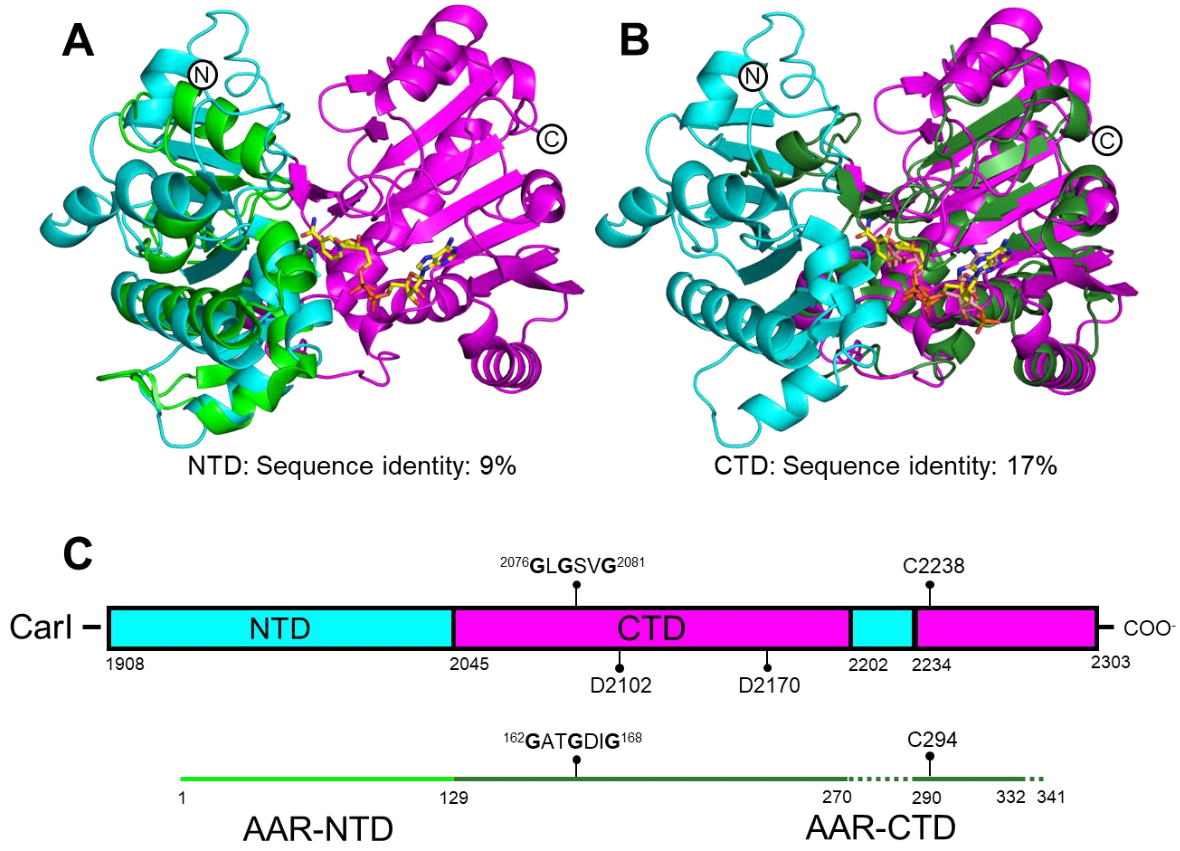


**Figure S13**. Comparison of the CarI TAD and *S. elongatus* PCC 7942 AAR (green, PDB 6JZY) structures. **A)** Superposition of N-terminal domains; RMSD = 3.2 Å for 79 Cα atoms. These domains have only 9% sequence identity. **B)** Superposition of C-terminal domains; RMSD = 1.6 Å for 120 Cα atoms. The $NAD^+$ (TAD, yellow) and NADPH (AAR, salmon) cofactors are similarly bound. The TAD and AAR CTDs are 19% identical. Compared to the NTDs, this higher identity is reflected in the greater structural similarity (3.2 Å vs. 1.6 Å RMSD). **C)** Schematic of TAD and AAR sequences. Regions of AAR that do not align structurally with TAD are indicated by dashed lines. AAR residues between 270 and 290 do not form an excursion into the NTD. The analogous conserved TAD Cys2238 and AAR Cys294 are indicated as well as the conserved GxGxxG “P loop” for cofactor phosphate binding. The conserved TAD Asp2102 and Asp2170 are not aspartates in AAR.

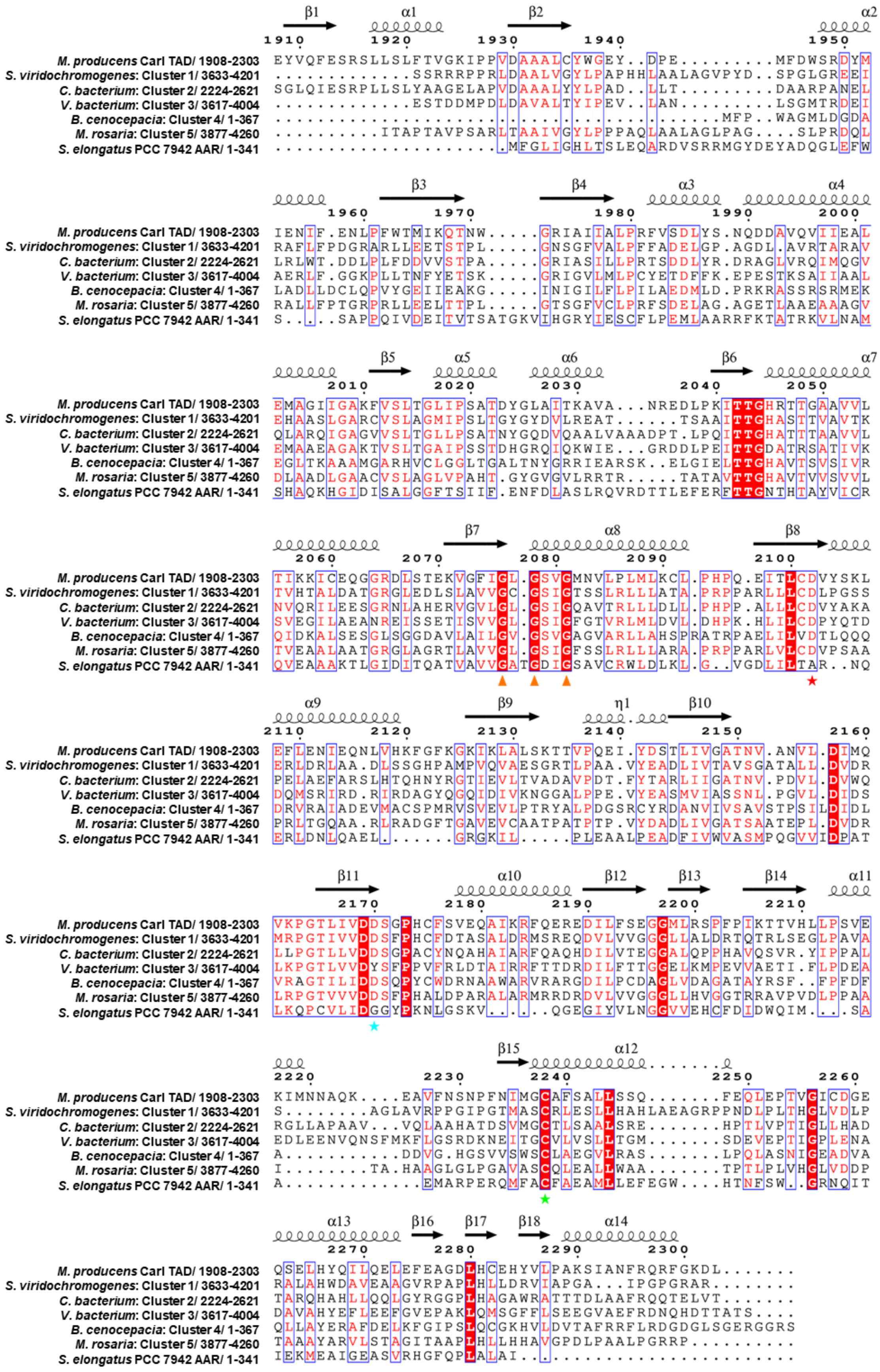
M. producens CarI TAD/ 1908-2303
S. viridochromogenes: Cluster 1/ 3633-4201
C. bacterium: Cluster 2/ 2224-2621
V. bacterium: Cluster 3/ 3617-4004
B. cenocepacia: Cluster 4/ 1-367
M. rosaria: Cluster 5/ 3877-4260
S. elongatus PCC 7942 AAR/ 1-341

**Figure S14.** Multiple sequence alignment of CarI TAD homologs, including the AAR. Representatives from each of the five largest SSN clusters were compared to the CarI TAD and *S. elongatus* PCC 7942 AAR[34]. Alignment was performed using Clustal Omega[21], followed by manual structure-based adjustment of the AAR alignment (PDB 6JZY) to the TADs. Secondary structure of CarI TAD is shown above. The η symbol marks a $3_{10}$-helix. Orange triangles mark the NAD binding site, a red star marks the phosphate-discriminating TAD Asp2102, a cyan star marks the Asp2170 that lies under the nicotinamide ring, and a green star highlights the putative catalytic Cys2238. Analysis as performed using ESPript 3.0[35]. Residue numbers of the aligned region are included in the row label. UniProt IDs: *Moorea producens* 3L CarI TAD: F4Y2B0; *Streptomyces viridichromogenes*: A0A0J7Z881; *Chloroflexacaea bacterium:* A0A968NAP9; *Verrucomicrobiales bacterium*: A0A2E0QPQ3; *Burkholderia cenocepacia*: A0A109EIG7; *Micromonospora rosaria:* A0A136PUW5; *Synechococcus elongatus* PCC 7942 AAR: Q54765.

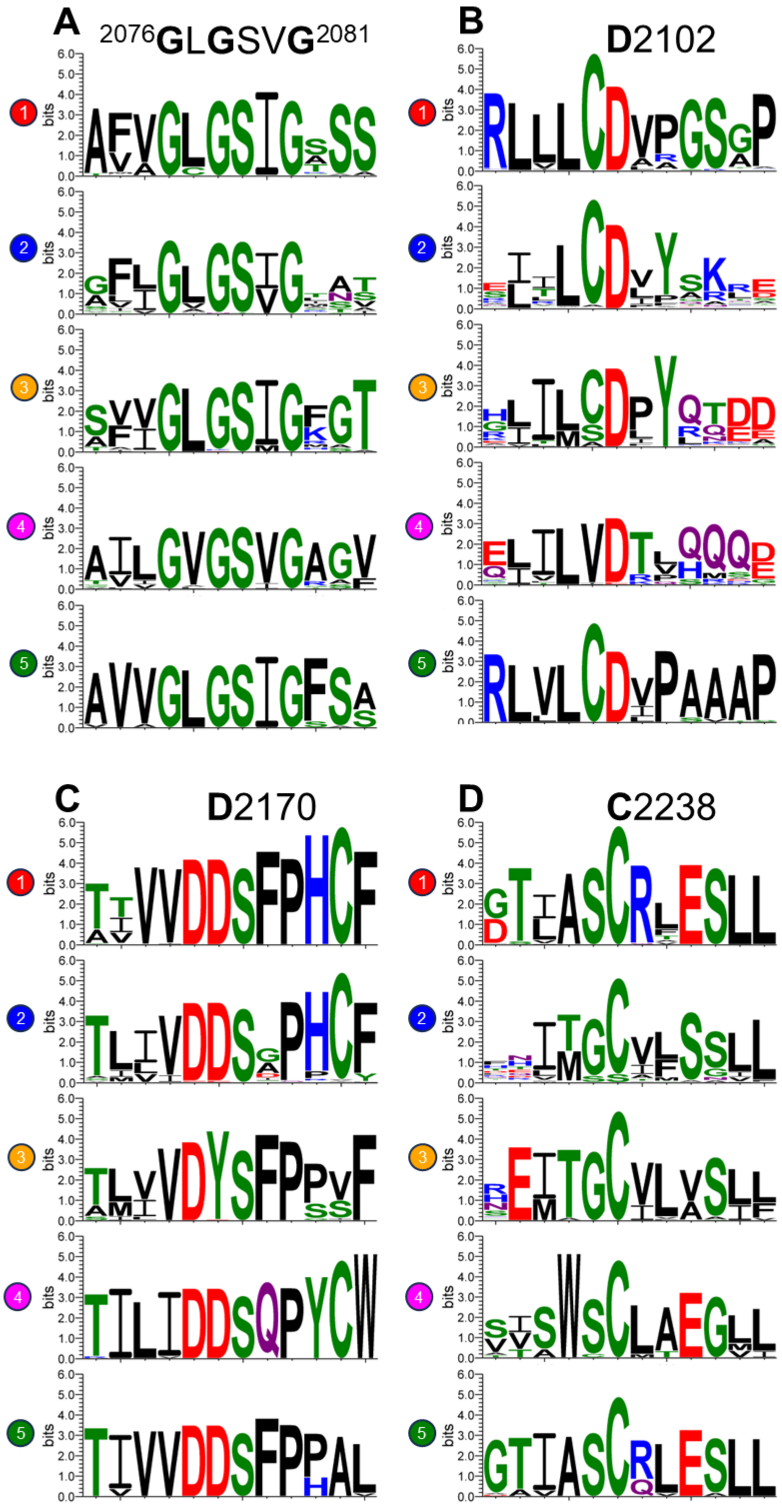


**Figure S15**. Stacked sequence logos for the consensus of clusters 1-5. The equivalent sequence position in CarI TAD is indicated at the top of each plot. **A)** The GXGXXG P-loop motif is intact in each case. **B)** Each TAD homolog selects against NADP(H) and CoA using the equivalent of Asp2102. **C)** CarI TAD Asp2170, which is positioned under the NAD nicotinamide ring, is conserved in clusters 1, 2, 4, and 5. In cluster 3, a tyrosine is instead present. **D)** Central Cys2238 is nearly invariant in each cluster. In cluster 2, this residue is rarely replaced by a serine that may perform similar function. Sequence logos were generated using WebLogo 3[36].

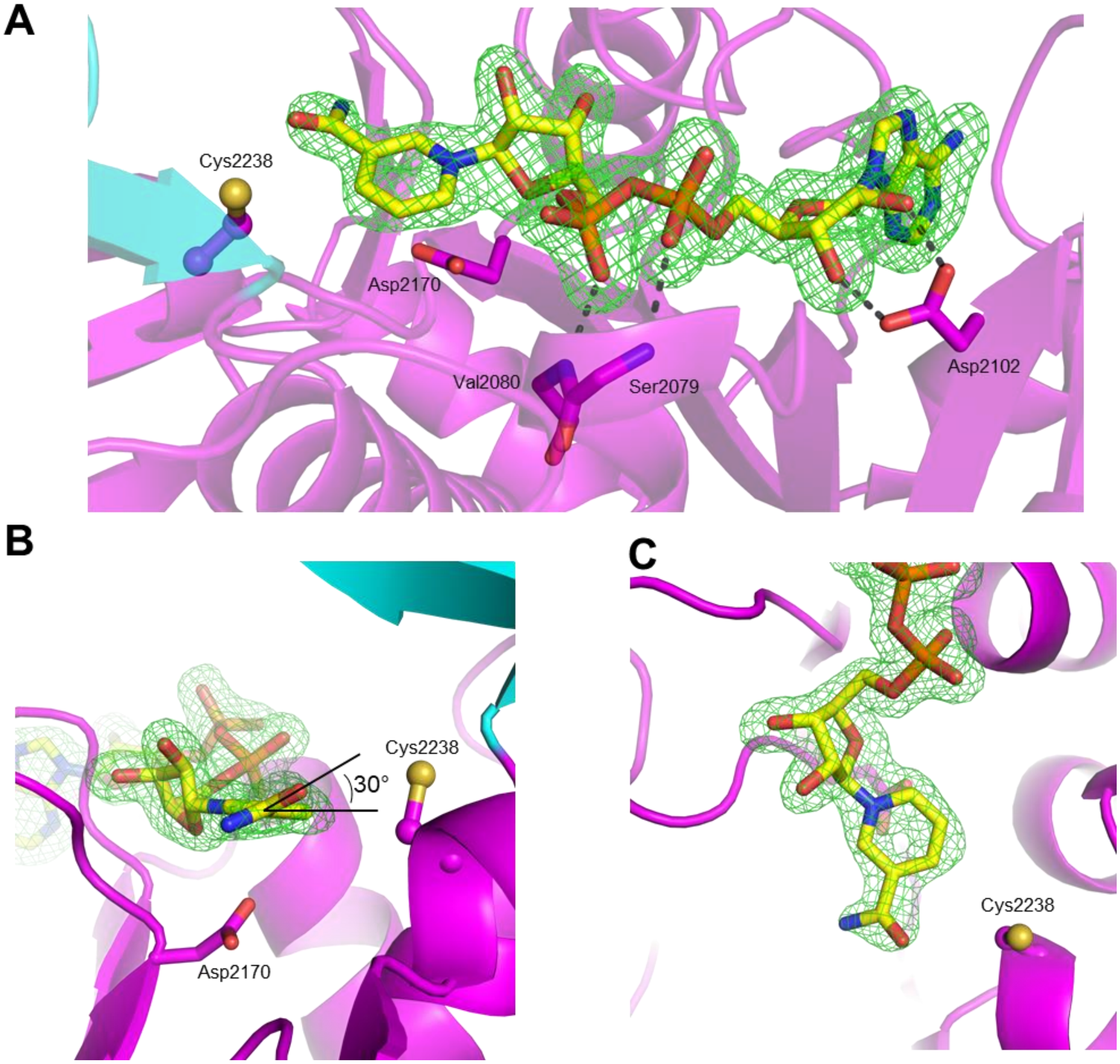


**Figure S16.** Details of NAD binding. NAD polder density[37] is contoured at 5σ. **A)** NAD is bound near CarI TAD Cys2238. Black dashed lines indicate important hydrogen bonds to NAD. Cys2238 is near redox-active nicotinamide C4 atom. Asp2102 forms selectivity-conferring hydrogen bonds with the 2' and 3' hydroxyl groups of NAD. **B**) The nicotinamide ring is well ordered despite the lack of hydrogen bonds. The amide plane is tipped 30° out of the aromatic ring plane. Conserved Asp2170 is nearby. **C**) View rotated 90°.

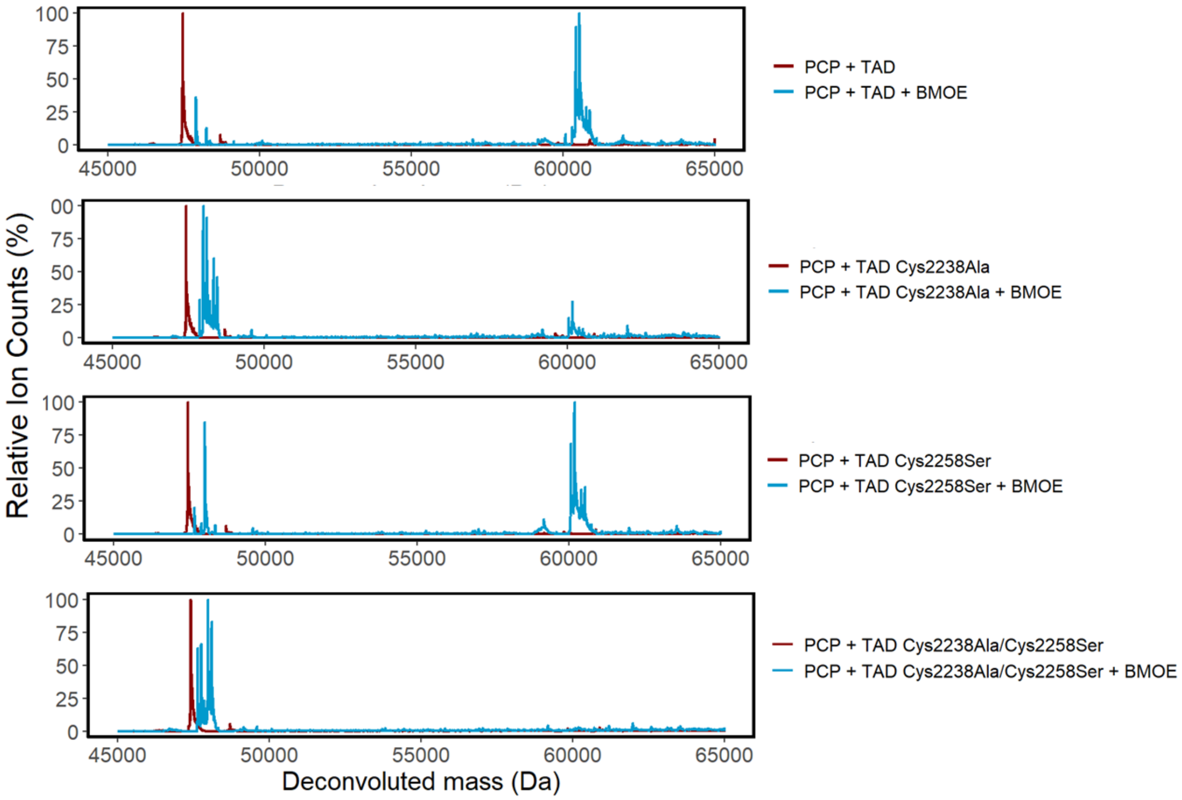


**Figure S17.** Deconvoluted mass spectrum of PCP-TAD crosslinking experiments with (blue) and without (red) BMOE. Mutagenesis of Cys2238 results in substantial decrease of PCP-TAD crosslink formation. Datasets are scaled to the highest peak of each spectrum.

**A**

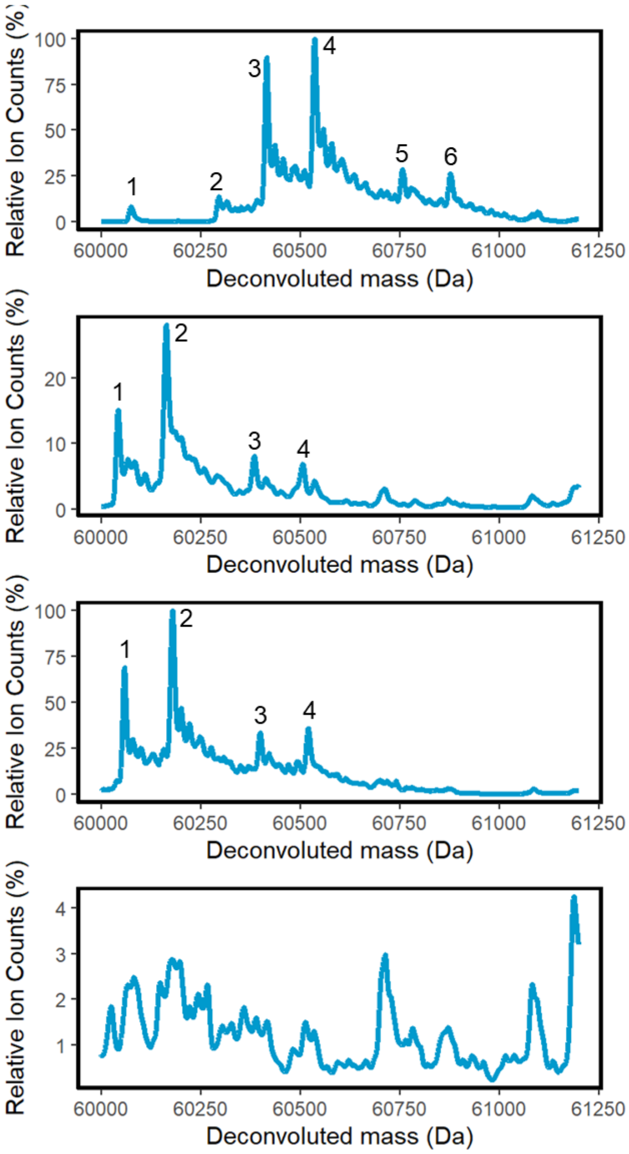


**B**

| PCP-BMOE-TAD + | Expected (Da) | Observed (Da) |
|---|---|---|
| (1) 1*BMOE | 60074.80 | 60074.95 |
| (2) 2*BMOE | 60294.99 | 60295.80 |
| (3) 1*BMOE, 1*BMOE-Cys | 60416.14 | 60416.25 |
| (4) 2*BMOE-Cys | 60537.29 | 60536.60 |
| (5) 2*BMOE-Cys, 1*BMOE | 60757.48 | 60757.15 |
| (6) 3*BMOE-Cys | 60878.63 | 60876.63 |

| PCP-BMOE-TAD C2238A + | Expected (Da) | Observed (Da) |
|---|---|---|
| (1) 1*BMOE | 60042.74 | 60042.90 |
| (2) 1*BMOE-Cys | 60163.90 | 60164.05 |
| (3) 1*BMOE, 1*BMOE-Cys | 60384.08 | 60384.80 |
| (4) 2*BMOE-Cys | 60505.23 | 60506.75 |

| PCP-BMOE-TAD C2238S + | Expected (Da) | Observed (Da) |
|---|---|---|
| (1) 1*BMOE | 60058.74 | 60059.00 |
| (2) 1*BMOE-Cys | 60178.89 | 60180.15 |
| (3) 1*BMOE, 1*BMOE-Cys | 60400.08 | 60399.80 |
| (4) 2*BMOE-Cys | 60521.23 | 60521.20 |

**PCP + TAD Cys2238Ala/ Cys2258Ser**

*No protein signal in this range*

**Figure S18.** Intact protein MS data for each crosslinking reaction. **A)** Deconvoluted mass spectrum in the PCP-TAD mass range. In addition to a PCP-TAD linkage via Cys2238, or less commonly Cys2258, up to three additional BMOE modification sites (with and without quenching by cysteine) can be detected. Datasets are scaled to the highest peak of each spectrum. **B)** Calculated and observed masses for each dominant peak.

A

B

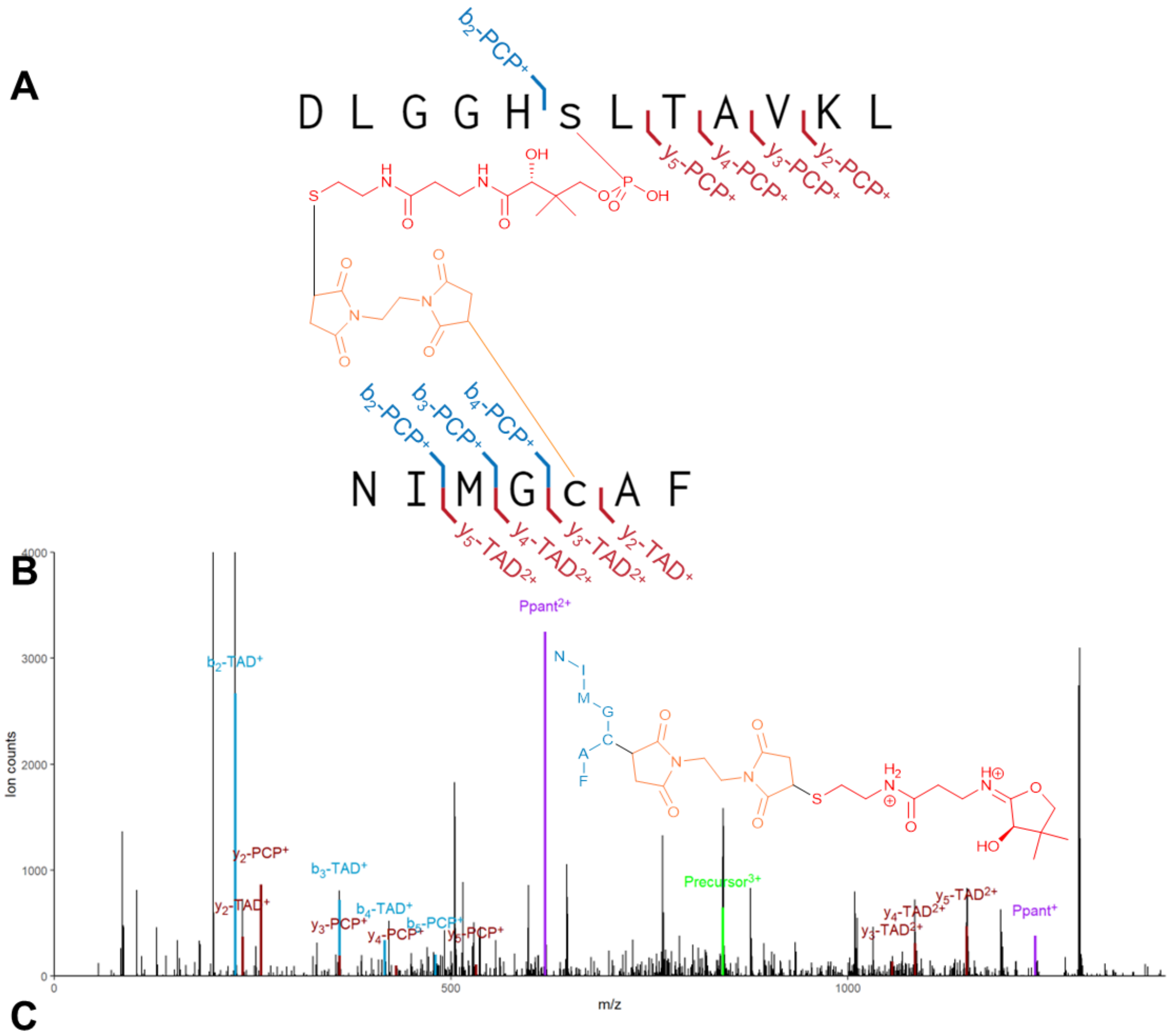


C

| Ion | Experimental Mass (*m/z*) | Theoretical Mass (*m/z*) | Mass error (ppm) |
|---|---|---|---|
| Precursor$^{3+}$ | 842.3809 | 842.3807 | 0.2374 |
| b2-TAD$^{+}$ | 228.1358 | 228.1343 | 6.5751 |
| y2-TAD$^{+}$ | 237.1247 | 237.1234 | 5.4824 |
| b3-TAD$^{+}$ | 359.1762 | 359.1747 | 4.1762 |
| b4-TAD$^{+}$ | 416.1967 | 416.1962 | 1.2014 |
| y3-TAD$^{2+}$ | 1055.4788 | 1055.4729 | 5.5899 |
| y4-TAD$^{2+}$ | 1083.9897 | 1083.9836 | 5.6274 |
| y5-TAD$^{2+}$ | 1149.4971 | 1149.5039 | -5.9156 |
| Ppant$^{+}$ | 1235.4863 | 1235.4894 | -2.5091 |
| Ppant$^{2+}$ | 618.253 | 618.2483 | 7.6021 |
| y2-PCP$^{+}$ | 260.1967 | 260.1969 | -0.7686 |
| y3-PCP$^{+}$ | 359.2654 | 359.2653 | 0.2783 |
| y4-PCP$^{+}$ | 430.3001 | 430.3024 | -5.3451 |
| b5-PCP$^{+}$ | 480.2233 | 480.2201 | 6.6636 |
| y5-PCP$^{+}$ | 531.3432 | 531.3501 | -12.9858 |

**Figure S19.** BMOE crosslinking of peptides containing the CarI PCP Ppant and TAD Cys2238. **A)** Schematic of peptides bridged by Ppant and BMOE groups. Interactive Peptide Spectral Annotator (IPSA)[38] was used to identify standard b/y ion matches. **B)** MS/MS spectrum resulting from fragmentation of the above triply charged species. A dominant Ppant ejection[39] peak is observed (purple). **C)** MS/MS statistics for this peptide.

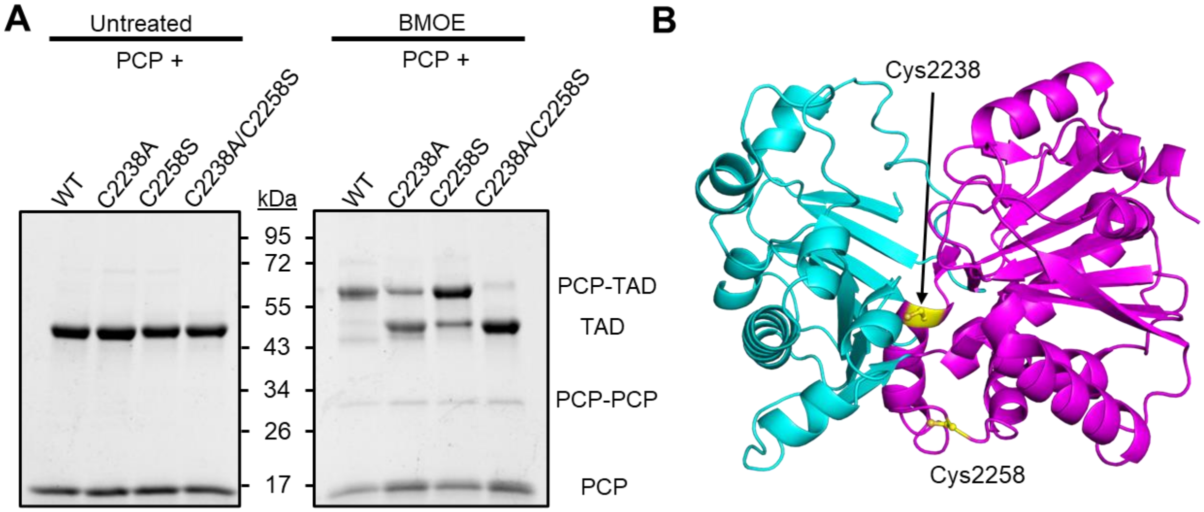


**Figure S20.** PCP-TAD crosslinking ability with mutagenized TAD proteins. **A)** SDS-PAGE gel of crosslinked PCP and mutagenized TADs. Both Cys2238 and Cys2258 participate in PCP crosslinking, but the effect from Cys2238 is dominant. **B)** Cartoon representation of the CarI TAD with both cysteines highlighted. Cys2258 is on the periphery of the protein and is an off-target crosslinking hit.

```
'''
Copyright (c) 2024 University of Michigan.  All rights reserved.

Developed by: Janet Smith Group
              University of Michigan
              https://www.lsi.umich.edu/science/our-labs/smith-lab

Permission is hereby granted, free of charge, to any person obtaining a
copy of this software and associated documentation files (the "Software"),
to deal with the Software without restriction, including without limitation
the rights to use, copy, modify, merge, publish, distribute, sublicense,
and/or sell copies of the Software, and to permit persons to whom the
Software is furnished to do so, subject to the following conditions:
* Redistributions of source code must retain the above copyright notice,
  this list of conditions and the following disclaimers.
* Redistributions in binary form must reproduce the above copyright notice,
  this list of conditions and the following disclaimers in the
  documentation and/or other materials provided with the distribution.
* Neither the names of Janet Smith Group, University of Michigan,
  nor the names of its contributors may be used to endorse or promote
  products derived from this Software without specific prior written
  permission.

THE SOFTWARE IS PROVIDED "AS IS", WITHOUT WARRANTY OF ANY KIND, EXPRESS OR
IMPLIED, INCLUDING BUT NOT LIMITED TO THE WARRANTIES OF MERCHANTABILITY,
FITNESS FOR A PARTICULAR PURPOSE AND NONINFRINGEMENT.  IN NO EVENT SHALL
THE CONTRIBUTORS OR COPYRIGHT HOLDERS BE LIABLE FOR ANY CLAIM, DAMAGES OR
OTHER LIABILITY, WHETHER IN AN ACTION OF CONTRACT, TORT OR OTHERWISE,
ARISING FROM, OUT OF OR IN CONNECTION WITH THE SOFTWARE OR THE USE OR OTHER
DEALINGS WITH THE SOFTWARE.
'''

import json       # Import json module for parsing JSON data
import argparse   # Import argparse for command-line argument parsing

# Function to parse command-line arguments
def parse_args():
    # Create an ArgumentParser object
    parser = argparse.ArgumentParser(
        description="Extract sequences from BLAST JSON output with length "
                    "requirement"
    )

    # Add arguments to the parser
    parser.add_argument(
        "input_file",
        help="Path to the input JSON file containing BLAST output"
    )
    parser.add_argument(
        "output_file",
        help="Path to the output file where results will be written"
    )
    parser.add_argument(
        "--min_length", type=int, default=1,
        help="Sequence length requirement (default: 1)"
    )

    # Return parsed arguments as a namespace object
    return parser.parse_args()

def main():
```

```
    # Parse arguments
    args = parse_args()

    # Try opening and reading input JSON file
    try:
        with open(args.input_file) as f:
            myjson = json.load(f)

    except IOError as e:
        print(f"Error opening or reading the input file: {e}")
        return

    except json.JSONDecodeError as e:
        print(f"Error decoding JSON from the input file: {e}")
        return

    # Try opening output file for writing
    try:
        with open(args.output_file, 'w') as ostr:
            # Iterate through each hit in JSON data
            for hit in
myjson["BlastOutput2"][0]["report"]["results"]["search"]["hits"]:
                descriptions = hit["description"]

                # Preferentially take an ID in UniProt format
                this_id = next(
                    (desc["id"].split('|')[1] for desc in descriptions
                     if desc["id"].startswith("gb|")),
                    descriptions[0]["id"].split('|')[1]
                )

                # Iterate through each high-scoring pair (HSP) in the hit
                for hitrange in hit["hsps"]:
                    sequence = hitrange["hseq"].replace('-', '')  # Remove gaps

                    if len(sequence) >= args.min_length:
                        # Only write sequences above minimum length
                        ostr.write(f">{this_id}\n{sequence}\n")

    except IOError as e:
        print(f"Error writing to the output file: {e}")
        return

# If this script is being run directly (rather than being imported), call main
if __name__ == "__main__":
    main()
```

**Figure S21**. Python script for the extraction of sequences that align to a domain of interest. This code accepts 3 input arguments: a BLAST results file in .json format, an output .txt file name, and an optional minimum length requirement. It returns a list of polypeptide IDs and sequences in FASTA format.

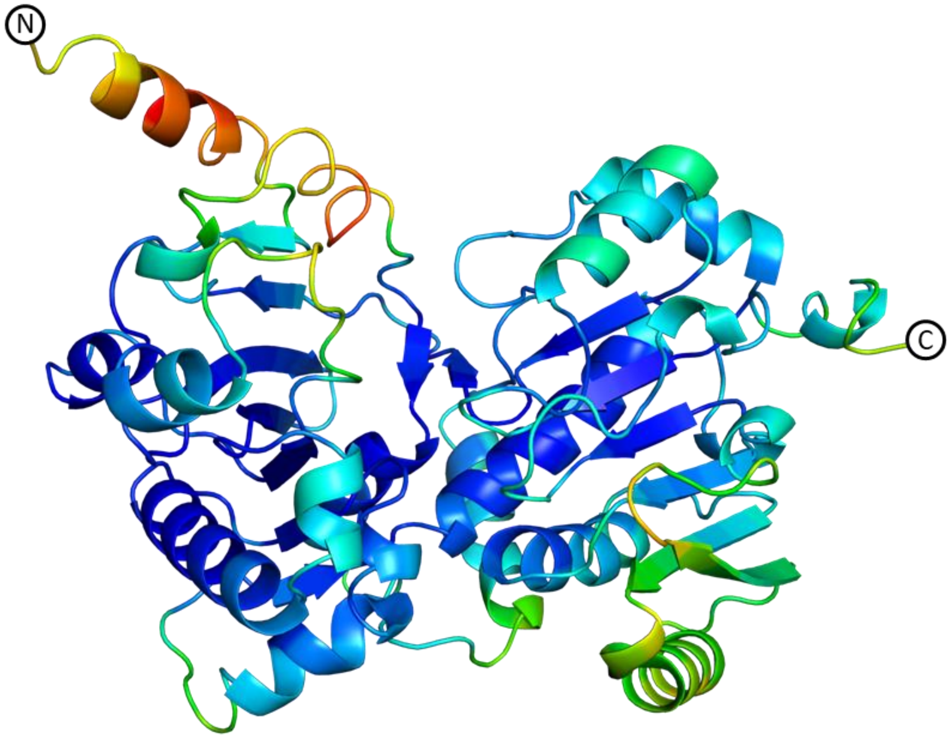


**Figure S22.** Chain A of the NADH-bound CarI TAD structure in space group $P3_221$ colored by B-factor. The N-terminal 20 residues pack poorly in the crystal lattice, reflected by the large B-factors. Removal of this helix resulted in a new crystal form and crystals of greater diffraction quality.

TABLES

**Table S1.** X-ray crystallography.

| | SeMet TAD | TAD | TAD:NADH | TAD | TAD:$NAD^+$ | TAD:dPCoA |
|---|---|---|---|---|---|---|
| Diffraction data | | | | | | |
| Space group | $P3_221$ | $P3_221$ | $P3_221$ | $P2_12_12_1$ | $P2_12_12_1$ | $P2_12_12_1$ |
| Unit cell a,b,c (Å) | 59.474, 59.474, 479.074 | 58.902, 58.902, 479.044 | 59.891, 59.891, 480.707 | 54.408, 74.549, 199.862 | 55.433, 72.813, 209.207 | 55.82, 72.867, 211.227 |
| X-ray source | APS 23ID-D | APS 23ID-B | APS 23ID-D | NSLS-II AMX | APS 23ID-D | ESRF ID30-B |
| Wavelength (Å) | 0.9794 | 1.0332 | 1.0332 | 0.9794 | 1.0332 | 0.8731 |
| $d_{min}$ (Å) | 3.37 (3.49-3.37)* | 2.28 (2.36-2.28) | 2.23 (2.31-2.23) | 2.82 (2.92-2.82) | 1.68 (1.74-1.68) | 1.94 (2.01-1.94) |
| $R_{merge}$ | 0.164 (0.932) | 0.071 (1.236) | 0.112 (1.624) | 0.164 (1.487) | 0.112 (1.498) | 0.092 (1.325) |
| Inner shell $R_{merge}$ | 0.063 | 0.044 | 0.064 | 0.058 | 0.057 | 0.048 |
| Mean I/σ(I) | 13.6 (2.5) | 14.0 (1.2) | 10.7 (0.9) | 10.9 (1.6) | 10.68 (1.15) | 11.2 (1.4) |
| Completeness (%) | 99.6 (95.8) | 99.88 (100.00) | 99.79 (99.96) | 94.63 (100.00) | 99.27 (98.53) | 99.8 (99.9) |
| Multiplicity | 11.4 (9.4) | 7.82 (7.83) | 9.17 (9.39) | 8.93 (9.54) | 7.7 (7.8) | 5.9 (6.2) |
| Total observations | 305,321 (24,948) | 359,494 (35,401) | 465,308 (46,643) | 172,368 (19,123) | 749,759 (73,254) | 379,436 (39,309) |
| Wilson B-factor (Å$^2$) | | 58.7 | 51.9 | 71.30 | 27.0 | 38.1 |
| $CC_{1/2}$ | 0.998 (0.864) | 0.999 (0.642) | 0.997 (0.713) | 0.997 (0.555) | 0.998 (0.512) | 0.997 (0.579) |
| Refinement | | | | | | |
| Data range (Å) | | 45.03-2.28 | 48.98-2.23 | 49.68-2.82 | 43.16-1.68 | 43.74-1.94 |
| Reflections in refinement (# (work / # free) | | 45,924 / 2,134 | 49,894 / 2,319 | 19,295 / 404 | 96,744 / 2,000 | 64,703 / 1,336 |
| $R_{work}/R_{free}$ (%) | | 0.235/0.267 | 0.220/0.263 | 0.239/0.300 | 0.202/0.229 | 0.198/0.224 |
| Non-hydrogen atoms (#) | | 6,560 | 6,595 | 5,983 | 6,828 | 6,510 |
| Protein | | 6,462 | 6,450 | 5,981 | 6,110 | 6,028 |
| Ligand | | 0 | 44 | 2 | 90 | 90 |
| Solvent | | 88 | 101 | 0 | 628 | 392 |
| Amino acids | | 821 | 819 | 765 | 777 | 773 |
| Deviation from ideality | | | | | | |
| Bond length (Å) | | 0.004 | 0.003 | 0.002 | 0.003 | 0.003 |
| Bond angle (°) | | 0.66 | 0.58 | 0.52 | 0.64 | 0.58 |
| Avg B-factor (Å$^2$) | | 78.4 | 68.2 | 73.3 | 35.7 | 45.7 |
| Protein | | 78.6 | 68.3 | 73.3 | 35.4 | 51.2 |
| Ligand | | NA | 73.9 | 89.9 | 27.3 | 42.11 |
| Solvent | | 66.0 | 59.0 | 0 | 39.5 | 46.9 |
| Ramachandran agreement | | | | | | |
| Favored (%) | | 94.85 | 97.17 | 96.83 | 97.92 | 97.49 |
| Allowed (%) | | 4.79 | 2.83 | 2.91 | 2.08 | 2.25 |
| Outlier (%) | | 0.36 | 0 | 0.26 | 0 | 0.26 |

* Values in parentheses correspond to the outermost shell of data.

**Table S2.** List of primers used in this study. Numbers indicate amino acid position with respect to the first methionine of *carI*. All sequences are listed 5' to 3'. Bolded nucleotides indicate restriction enzyme cut sites or LIC handles. Red nucleotides indicate mutated sites.

| Primer | Sequence |
|---|---|
| CarI Module 1 NdeI F | **CATATG**AAAACAGTTAATTTT |
| CarI Module 2303 XhoI R | **CTCGAG**CAAATCTTTGCCAAA |
| CarI TAD 1888 NdeI F | **CATATG**TCAATTAATAATGAA |
| CarI TAD Δhelix 1908 LIC F | **TACTTCCAATCCAATGCC**GAGTATGTACAATTT |
| CarI TAD 2303 LIC R | **TTATCCACTTCCAATG**TTACAAATCTTTGCC |
| CarI PCP 1790 NdeI F | **CATATG**GTGGCCAGTAGTGTG |
| CarI PCP 1887 XhoI R | **CTCGAG**TTGAGAAATTGGCTC |
| CarI TAD Cys2238Ala F | AGCAGTTTTTAATTCAAATCCTTTTAATATCATGGGTGCTGCTTTTTCTGCTTTACTAT |
| CarI TAD Cys2238Ala R | ATAGTAAAGCAGAAAAAGCAGCACCCATGATATTAAAAGGATTTGAATTAAAAACTGCT |
| CarI TAD Cys2258Ser F | ACTGGAGCCTACTGTAGGAATTAGCGATGGAGAAC |
| CarI TAD Cys2258Ser R | GTTCTCCATCGCTAATTCCTACAGTAGGCTCCAGT |

**Table S3.** List of constructs used in this study. Numbers indicate amino acid position with respect to CarI Met1.

| Name | Vector | Insert | Construct Design |
|---|---|---|---|
| CarI TAD | pET29b(+) | CarI TAD: 1888 - 2303 | 6xHis - Gene |
| CarI PCP | pET29b(+) | CarI PCP: 1790 - 1887 | 6xHis - Gene |
| pMocr: PCP-TAD | pMocr | CarI PCP-TAD: 1790 - 2303 | 6xHis - Mocr - TEV - Gene |
| pMR88 | pMCSG7 | CarI TAD Δhelix: 1908 - 2303 | 6xHis - TEV - Gene |
| pMR79 | pET29b(+) | CarI TAD C2238A: 1888 - 2303 | 6xHis - Gene |
| pMR113 | pET29b(+) | CarI TAD C2258S: 1888 - 2303 | 6xHis - Gene |
| pMR114 | pET29b(+) | CarI TAD C2238A/C2258S: 1888 - 2303 | 6xHis - Gene |

SUPPLEMENTAL INFORMATION REFERENCES

(14) Liebschner, D.; Afonine, P. V.; Baker, M. L.; Bunkóczi, G.; Chen, V. B.; Croll, T. I.; Hintze, B.; Hung, L.-W.; Jain, S.; McCoy, A. J.; Moriarty, N. W.; Oeffner, R. D.; Poon, B. K.; Prisant, M. G.; Read, R. J.; Richardson, J. S.; Richardson, D. C.; Sammito, M. D.; Sobolev, O. V.; Stockwell, D. H.; Terwilliger, T. C.; Urzhumtsev, A. G.; Videau, L. L.; Williams, C. J.; Adams, P. D. Macromolecular Structure Determination Using X-Rays, Neutrons and Electrons:

Recent Developments in Phenix. *Acta Crystallogr. Sect. Struct. Biol.* **2019**, *75* (10), 861–877. https://doi.org/10.1107/S2059798319011471.
(15) Cowtan, K. The Buccaneer Software for Automated Model Building. 1. Tracing Protein Chains. *Acta Crystallogr. D Biol. Crystallogr.* **2006**, *62* (9), 1002–1011. https://doi.org/10.1107/S0907444906022116.
(16) Emsley, P.; Lohkamp, B.; Scott, W. G.; Cowtan, K. Features and Development of *Coot*. *Acta Crystallogr. D Biol. Crystallogr.* **2010**, *66* (4), 486–501. https://doi.org/10.1107/S0907444910007493.
(17) Afonine, P. V.; Grosse-Kunstleve, R. W.; Echols, N.; Headd, J. J.; Moriarty, N. W.; Mustyakimov, M.; Terwilliger, T. C.; Urzhumtsev, A.; Zwart, P. H.; Adams, P. D. Towards Automated Crystallographic Structure Refinement with Phenix.Refine. *Acta Crystallogr. D Biol. Crystallogr.* **2012**, *68* (4), 352–367. https://doi.org/10.1107/S0907444912001308.
(18) McCoy, A. J.; Grosse-Kunstleve, R. W.; Adams, P. D.; Winn, M. D.; Storoni, L. C.; Read, R. J. Phaser Crystallographic Software. *J. Appl. Crystallogr.* **2007**, *40* (4), 658–674. https://doi.org/10.1107/S0021889807021206.
(19) Schrödinger, LLC. The PyMOL Molecular Graphics System, Version 1.8, 2015.
(20) Chen, V. B.; Arendall, W. B.; Headd, J. J.; Keedy, D. A.; Immormino, R. M.; Kapral, G. J.; Murray, L. W.; Richardson, J. S.; Richardson, D. C. MolProbity: All-Atom Structure Validation for Macromolecular Crystallography. *Acta Crystallogr. D Biol. Crystallogr.* **2010**, *66* (Pt 1), 12–21. https://doi.org/10.1107/S0907444909042073.
(21) Sievers, F.; Higgins, D. G. Clustal Omega for Making Accurate Alignments of Many Protein Sequences. *Protein Sci. Publ. Protein Soc.* **2018**, *27* (1), 135–145. https://doi.org/10.1002/pro.3290.
(22) Waterhouse, A. M.; Procter, J. B.; Martin, D. M. A.; Clamp, M.; Barton, G. J. Jalview Version 2—a Multiple Sequence Alignment Editor and Analysis Workbench. *Bioinformatics* **2009**, *25* (9), 1189–1191. https://doi.org/10.1093/bioinformatics/btp033.
(23) Fraczkiewicz, R.; Braun, W. Exact and Efficient Analytical Calculation of the Accessible Surface Areas and Their Gradients for Macromolecules. *J. Comput. Chem. 19* (3).
(24) Gedgaudas, M.; Baronas, D.; Kazlauskas, E.; Petrauskas, V.; Matulis, D. Thermott: A Comprehensive Online Tool for Protein–Ligand Binding Constant Determination. *Drug Discov. Today* **2022**, *27* (8), 2076–2079. https://doi.org/10.1016/j.drudis.2022.05.008.
(25) Engene, N.; Rottacker, E. C.; Kaštovský, J.; Byrum, T.; Choi, H.; Ellisman, M. H.; Komárek, J.; Gerwick, W. H. Moorea Producens Gen. Nov., Sp. Nov. and Moorea Bouillonii Comb. Nov., Tropical Marine Cyanobacteria Rich in Bioactive Secondary Metabolites. *Int. J. Syst. Evol. Microbiol.* **2012**, *62* (5), 1171–1178. https://doi.org/10.1099/ijs.0.033761-0.
(26) Letunic, I.; Bork, P. Interactive Tree of Life (iTOL) v6: Recent Updates to the Phylogenetic Tree Display and Annotation Tool. *Nucleic Acids Res.* **2024**, *52* (W1), W78–W82. https://doi.org/10.1093/nar/gkae268.
(27) Blin, K.; Shaw, S.; Steinke, K.; Villebro, R.; Ziemert, N.; Lee, S. Y.; Medema, M. H.; Weber, T. antiSMASH 5.0: Updates to the Secondary Metabolite Genome Mining Pipeline. *Nucleic Acids Res.* **2019**, *47* (W1), W81–W87. https://doi.org/10.1093/nar/gkz310.
(28) Sanchez, L. M.; Lopez, D.; Vesely, B. A.; Togna, G. D.; Gerwick, W. H.; Kyle, D. E.; Linington, R. G. Almiramides A–C: Discovery and Development of a New Class of Leishmaniasis Lead Compounds. *J. Med. Chem.* **2010**, *53* (10), 4187–4197. https://doi.org/10.1021/jm100265s.
(29) Maloney, F. P.; Gerwick, L.; Gerwick, W. H.; Sherman, D. H.; Smith, J. L. Anatomy of the β-Branching Enzyme of Polyketide Biosynthesis and Its Interaction with an Acyl-ACP Substrate. *Proc. Natl. Acad. Sci. U. S. A.* **2016**, *113* (37), 10316–10321. https://doi.org/10.1073/pnas.1607210113.
(30) Reitz, Z. L.; Hardy, C. D.; Suk, J.; Bouvet, J.; Butler, A. Genomic Analysis of Siderophore β-Hydroxylases Reveals Divergent Stereocontrol and Expands the Condensation Domain

Family. *Proc. Natl. Acad. Sci.* **2019**, *116* (40), 19805–19814. https://doi.org/10.1073/pnas.1903161116.
(31) Hutchinson, E. G.; Thornton, J. M. PROMOTIF--a Program to Identify and Analyze Structural Motifs in Proteins. *Protein Sci. Publ. Protein Soc.* **1996**, *5* (2), 212–220.
(32) Laskowski, R. A.; Hutchinson, E. G.; Michie, A. D.; Wallace, A. C.; Jones, M. L.; Thornton, J. M. PDBsum: A Web-Based Database of Summaries and Analyses of All PDB Structures. *Trends Biochem. Sci.* **1997**, *22* (12), 488–490. https://doi.org/10.1016/S0968-0004(97)01140-7.
(33) Bond, C. S. TopDraw: A Sketchpad for Protein Structure Topology Cartoons. *Bioinforma. Oxf. Engl.* **2003**, *19* (2), 311–312. https://doi.org/10.1093/bioinformatics/19.2.311.
(34) Gao, Y.; Zhang, H.; Fan, M.; Jia, C.; Shi, L.; Pan, X.; Cao, P.; Zhao, X.; Chang, W.; Li, M. Structural Insights into Catalytic Mechanism and Product Delivery of Cyanobacterial Acyl-Acyl Carrier Protein Reductase. *Nat. Commun.* **2020**, *11* (1), 1–11. https://doi.org/10.1038/s41467-020-15268-y.
(35) Robert, X.; Gouet, P. Deciphering Key Features in Protein Structures with the New ENDscript Server. *Nucleic Acids Res.* **2014**, *42* (W1), W320–W324. https://doi.org/10.1093/nar/gku316.
(36) Crooks, G. E.; Hon, G.; Chandonia, J.-M.; Brenner, S. E. WebLogo: A Sequence Logo Generator. *Genome Res.* **2004**, *14* (6), 1188–1190. https://doi.org/10.1101/gr.849004.
(37) Liebschner, D.; Afonine, P. V.; Moriarty, N. W.; Poon, B. K.; Sobolev, O. V.; Terwilliger, T. C.; Adams, P. D. Polder Maps: Improving OMIT Maps by Excluding Bulk Solvent. *Acta Crystallogr. Sect. Struct. Biol.* **2017**, *73* (2), 148–157. https://doi.org/10.1107/S2059798316018210.
(38) Brademan, D. R.; Riley, N. M.; Kwiecien, N. W.; Coon, J. J. Interactive Peptide Spectral Annotator: A Versatile Web-Based Tool for Proteomic Applications. *Mol. Cell. Proteomics MCP* **2019**, *18* (8 suppl 1), S193–S201. https://doi.org/10.1074/mcp.TIR118.001209.
(39) Dorrestein, P. C.; Bumpus, S. B.; Calderone, C. T.; Garneau-Tsodikova, S.; Aron, Z. D.; Straight, P. D.; Kolter, R.; Walsh, C. T.; Kelleher, N. L. Facile Detection of Acyl and Peptidyl Intermediates on Thiotemplate Carrier Domains via Phosphopantetheinyl Elimination Reactions during Tandem Mass Spectrometry. *Biochemistry* **2006**, *45* (42), 12756–12766. https://doi.org/10.1021/bi061169d.
(40) Hooper, G. J.; Orjala, J.; Schatzman, R. C.; Gerwick, W. H. Carmabins A and B, New Lipopeptides from the Caribbean Cyanobacterium *Lyngbya Majuscula*. *J. Nat. Prod.* **1998**, *61* (4), 529–533. https://doi.org/10.1021/np970443p.
(41) Jones, A. C.; Monroe, E. A.; Podell, S.; Hess, W. R.; Klages, S.; Esquenazi, E.; Niessen, S.; Hoover, H.; Rothmann, M.; Lasken, R. S.; Yates, J. R.; Reinhardt, R.; Kube, M.; Burkart, M. D.; Allen, E. E.; Dorrestein, P. C.; Gerwick, W. H.; Gerwick, L. Genomic Insights into the Physiology and Ecology of the Marine Filamentous Cyanobacterium Lyngbya Majuscula. *Proc. Natl. Acad. Sci. U. S. A.* **2011**, *108* (21), 8815–8820. https://doi.org/10.1073/pnas.1101137108.
(42) Leao, T.; Castelão, G.; Korobeynikov, A.; Monroe, E. A.; Podell, S.; Glukhov, E.; Allen, E. E.; Gerwick, W. H.; Gerwick, L. Comparative Genomics Uncovers the Prolific and Distinctive Metabolic Potential of the Cyanobacterial Genus Moorea. *Proc. Natl. Acad. Sci. U. S. A.* **2017**, *114* (12), 3198–3203. https://doi.org/10.1073/pnas.1618556114.
(43) Jiménez, J. I.; Scheuer, P. J. New Lipopeptides from the Caribbean Cyanobacterium Lyngbya Majuscula. *J. Nat. Prod.* **2001**, *64* (2), 200–203. https://doi.org/10.1021/np000462q.
(44) McPhail, K. L.; Correa, J.; Linington, R. G.; González, J.; Ortega-Barría, E.; Capson, T. L.; Gerwick, W. H. Antimalarial Linear Lipopeptides from a Panamanian Strain of the Marine Cyanobacterium Lyngbya Majuscula. *J. Nat. Prod.* **2007**, *70* (6), 984–988. https://doi.org/10.1021/np0700772.

(45) Moss, N. A.; Seiler, G.; Leão, T. F.; Castro-falcón, G.; Gerwick, L.; Chambers, C.; Gerwick, W. H. Combinatorial Assembly-Line Biosynthesis Produces Vatiamides A-F. *Angew. Chem. Int. Ed.* **2019**, *58* (27), 9027–9031.
(46) Ngo, T.-E.; Ecker, A.; Ryu, B.; Guild, A.; Remmel, A.; Boudreau, P. D.; Alexander, K. L.; Naman, C. B.; Glukhov, E.; Avalon, N. E.; Shende, V. V.; Thomas, L.; Dahesh, S.; Nizet, V.; Gerwick, L.; Gerwick, W. H. Structure and Biosynthesis of Hectoramide B, a Linear Depsipeptide from Marine Cyanobacterium Moorena Producens JHB Discovered via Coculture with Candida Albicans. *ACS Chem. Biol.* **2024**, *19* (3), 619–628. https://doi.org/10.1021/acschembio.3c00391.
(47) Marner, F. J.; Moore, R. E.; Hirotsu, K.; Clardy, J. Majusculamides A and B, Two Epimeric Lipodipeptides from Lyngbya Majuscula Gomont. *J. Org. Chem.* **1977**, *42* (17), 2815–2819. https://doi.org/10.1021/jo00437a005.
(48) Bracegirdle, J.; Casandra, D.; Rocca, J. R.; Adams, J. H.; Baker, B. J. Highly N-Methylated Peptides from the Antarctic Sponge Inflatella Coelosphaeroides Are Active against Plasmodium Falciparum. *J. Nat. Prod.* **2022**, *85* (10), 2454–2460. https://doi.org/10.1021/acs.jnatprod.2c00684.
